# Limited predictability of tree-level responses to drought across European forests

**DOI:** 10.64898/2026.02.26.708208

**Authors:** Diego I. Rodríguez-Hernández, Fabian J. Fischer, Duncan A. O’Brien, Martin De Kauwe, Bo Wang, Olivier Bouriaud, Tommaso Jucker

**Author notes:** **Corresponding author**: Diego I. Rodríguez-Hernández.

## Abstract

Climate change is increasing the frequency, duration and severity of extreme events such as heatwaves and droughts, pushing trees near or beyond their ecophysiological limits. Understanding what governs variability in how trees respond to drought – such as intrinsic factors related to their size, age, and species, or extrinsic factors shaped by their local competitive environment – is critical for predicting long-term forest resilience to climate change and developing climate-smart forest management strategies. Here, we use tree ring data from 2909 trees belonging to sixteen species distributed across Europe’s major forest types to comprehensively assess what factors contribute most to a tree’s ability to withstand and recover from extreme drought events. We found that trees with larger living crowns generally exhibited higher post-drought growth recovery and resilience, while trees exposed to lower drought intensities showed greater resistance. Conversely, neither the density nor the diversity of a tree’s local competitive neighbourhood had any clear influence on its response to drought. More generally, we found that our ability to predict whether a tree would exhibit resilience to drought was low (*R*^2^ = 13–21) and was largely driven by species-specific responses and topographic variation across forest types, rather than by tree- and stand-level attributes. These findings highlight that drought responses are inherently complex and strongly influenced by forest type and by heterogeneous responses among species. Integrating tree-ring, physiological, and remote-sensing data with mechanistic models represents a promising avenue for improving forecasts of future forest resilience to climate change.

## INTRODUCTION

Climate change is projected to make summer extremes more frequent, longer-lasting, and more severe. Consequently, forest ecosystems worldwide face increasing heat and drought-induced tree mortality risks (Allen et al., 2010; Hammond et al., 2022; McDowell et al., 2018, 2022). This situation has led to declining crown health, reduced tree growth and survival rates, and widespread forest dieback in recent decades (Camarero et al., 2024; Liu et al., 2023; Sterck et al., 2024). Given that the future of forests remains uncertain (Hammond et al., 2022; Sterck et al., 2024), improving our understanding of their resilience to rapid climate change is critical (Haberstroh & Werner, 2022). In this regard, although a significant amount of research has enhanced our understanding of how tree species respond to drought (Anderegg et al., 2015; Gazol et al., 2018; McDowell et al., 2022), most studies have focused on explaining differences among species groups (Castagneri et al., 2022; Pardos et al., 2021; Song, Sterck, Sass Klaassen, et al., 2022; Song, Sterck, Zhou, et al., 2022), often with small sample sizes or limited to a restricted set of tree species and forest types (Bottero et al., 2021; Gazol & Camarero, 2016; Schmied, Hilmers, et al., 2023; Schmied, Pretzsch, et al., 2023; Song, Sterck, Sass Klaassen, et al., 2022; Zang et al., 2012, 2014). This knowledge gap hinders our ability to forecast local-scale effects of drought across a larger set of tree species and forest types and limits the extent to which we can translate ecological knowledge into landscape-scale forest management interventions aimed at enhancing forest ecosystem resilience to climate change.

There are several reasons why some trees might be more resilient than others. This includes factors intrinsic to the tree itself, such as its age, size, crown dimensions, and root investment. Several studies have demonstrated that older trees with a taller stature are more prone to hydraulic failure (Bennett et al., 2015; Stovall et al., 2019). This increased vulnerability is hypothesised to arise from the longer distance water travels from the roots to the canopy (Fernández de Uña et al., 2023; Ma et al., 2023), along with slower growth rates and reduced photosynthate availability, limiting investment in defence and new tissue growth. Yet, the role of height during drought remains unresolved (Ma et al., 2023), given that it has only been tested at the species level (but see Fernández de Uña et al., 2023; González de Andrés et al., 2021). For example, taller trees might also be more resilient to drought as they usually have greater investment in roots and greater trunk water capacitance (Fernández de Uña et al., 2023; González de Andrés et al., 2021). Other aspects of tree architecture related to crown dimensions could also have a beneficial impact on drought tolerance and recovery, such as having deeper and wider crowns that enhance self-shading and help minimise water loss via transpiration (Ma et al., 2023; Su et al., 2020). Moreover, a tree’s pre-drought growth conditions (i.e., mean growth rate) may also influence its ability to withstand and recover in its aftermath, with faster growth rates often associated with lower resilience (Bose et al., 2020; Martínez-Vilalta et al., 2012; Tao et al., 2024; Zang et al., 2014).

Other extrinsic (i.e., external) factors can also affect a tree’s resilience to drought. For instance, the frequency, severity, and duration of the drought itself can have a significant influence on whether trees can sustain growth during and after drought (Bose et al., 2020; Gao et al., 2018; Serrano León et al., 2025; Zhang et al., 2022). Equally, a tree’s local competitive environment can shape its response to drought. For example, trees growing in drier and more densely packed stands may be more prone to drought stress and hydraulic failure due to intensified competition for water and higher evaporative demands (Haberstroh & Werner, 2022; Ma et al., 2023; Young et al., 2017). These competitive effects may be mitigated – at least in part – when trees grow in structurally more complex communities (Ma et al., 2023) or alongside functionally complementary neighbours that reduce water competition or promote facilitative interactions during moderate drought stress (Grossiord, 2020; Haberstroh & Werner, 2022; Serrano León et al., 2025). However, such benefits can diminish or reverse under extreme conditions (Haberstroh & Werner, 2022; Serrano León et al., 2025). Species mixing has been hypothesised to reduce tree-to-tree competition via niche partitioning and facilitative interactions, potentially lessening the impacts of drought (Pardos et al., 2021; Pretzsch et al., 2013). However, while there is robust evidence that mixed-species forests are on average more productive than monocultures (del Río et al., 2022; Jucker, Bouriaud, Avacaritei, & Coomes, 2014), support for positive diversity effects on drought resilience is less conclusive, with studies reporting contrasting outcomes depending on the mixture of tree species, their functional traits, mycorrhizal type, and the size structure of the community (Forrester et al., 2016; Grossiord, 2020; Grossiord, Granier, Ratcliffe, et al., 2014; Pardos et al., 2021; Sachsenmaier et al., 2024; Serrano León et al., 2025).

To comprehensively assess what factors govern how trees respond to drought, we used tree-ring records to reconstruct tree-level growth responses across Europe’s major forest types during extreme drought events. In total, we sampled tree cores from 2909 trees spanning a gradient of 20° in latitude from Mediterranean forests in Spain and Italy to boreal forests in Finland. Tree cores were acquired from 16 tree species that include all of Europe’s dominant coniferous, deciduous broadleaf, and evergreen broadleaf trees. We coupled these individual growth records with data on a range of intrinsic and extrinsic factors believed to play a role in shaping how trees respond to drought, including information on tree height, age, crown dimensions, growth conditions prior to drought and local competitive environment. Leveraging this unique dataset, we sought to address the following questions that motivated our research: (1) How does resilience to drought vary among major European tree species and forest types? (2) What are the primary factors that shape a tree’s resilience to drought? And (3) How predictable is individual tree-level resilience to drought?

## METHODS

### Forest types and species

Research was conducted across the FunDivEUROPE forest plot network, comprising 209 permanent plots distributed across six forest types in Europe. The plot network was originally established in 2011 to test for the effects of tree diversity on ecosystem functioning in natural forests across Europe (Baeten et al., 2013), but also provides an ideal platform to evaluate forest responses to climate change on a continental scale (Grossiord, Granier, Ratcliffe, et al., 2014; Jucker et al., 2016, 2017). The six forest types span more than 20° in latitude and encompass the major forest ecosystems found in Europe, including boreal forests in Finland, hemi-boreal and mixed broadleaved-coniferous forests in Poland, beech forests in Germany, mountainous beech forests in Romania, thermophilous deciduous forests in Italy and Mediterranean mixed forests in Spain (Table S1; Baeten et al., 2013). Plots were established in mature stands at mid-to-late stages of stem exclusion, except in Finland, where stands are even-aged and have regenerated following clear-cutting that occurred around 50 years prior (Baeten et al., 2013; Jucker, Bouriaud, Avacaritei, & Coomes, 2014). The plot network captures 16 dominant European tree species, several of which are found at more than one forest type (e.g., *Picea abies*, *Pinus sylvestris* and *Fagus sylvatica*). The species pool includes four coniferous, eleven deciduous broadleaves and one evergreen broadleaf species (see Table S1 for a full species list).

In 2011, 209 permanent plots (30 × 30 m in size) were established across the network in stands with varying levels of tree diversity – from pure monocultures to mixtures of multiple species (Baeten et al., 2013). Within each plot, all stems with a trunk diameter ≥7.5 cm were measured, mapped, identified to species, and permanently tagged, resulting in a total of 12,939 measured stems across the network (Jucker et al., 2017). For each stem, the diameter at breast height (DBH, cm) was recorded using a diameter tape, while tree height (H, m), crown radius (CR, m; as the average of two orthogonal measurements) and crown depth (CD, m) were measured using a vertex hypsometer (Jucker et al., 2015).

### Wood coring and dendrochronological analysis

In 2012, we collected increment cores from 2909 trees across 208 plots of the network following a size-stratified random sampling approach (Jucker et al., 2016, 2017). Wood core sampling and processing are described in detail in Jucker et al. (2014). Briefly, wood cores were collected at breast height using a 5.15-mm-diameter Haglöf increment borer. Once collected, cores were mounted, sanded and imaged using a high-resolution flatbed scanner (2400 dpi). Annual ring width measurements and cross-dating were then performed using the CDendro software suite (Cybis Elektronik & Data, Saltsjöbaden, Sweden). Yearly radial growth increments (mm yr^−1^) were subsequently converted to basal area increments (BAI; cm^2^ yr^−1^) by assuming circular stems. To remove ontogenetic growth trends from the BAI timeseries and facilitate the detection of drought years (Klesse & Bigler, 2025), we detrended all BAI timeseries (hereafter BAI_detrend_; unitless) using Friedman’s super smoother spline method as implemented (Fig. S1; Pardos et al., 2021) in the *dplR* package in R (Bunn, 2008). For summary statistics relating to cross-dating and temporal sample coverage, see Table S2.

### Identifying drought events

To identify site-specific drought events responsible for growth reductions, we used climatic water deficit (CWD) during the growing season (April to September) as an indicator of plant water stress (Anderegg et al., 2015; Hammond et al., 2022; Schwarz et al., 2020). CWD combines the effects of solar radiation, potential evapotranspiration, precipitation, and air temperature on watershed conditions, given a water-balance estimate of available soil moisture (Stephenson, 1998). Monthly CWD values were obtained for each plot from the TerraClimate database, which covers the period from 1958 to 2017 at a spatial resolution of ∼4 km^2^ (Abatzoglou et al., 2018). Although this resolution does not capture fine-scale microclimatic variation, it adequately represents plot-level water deficits and soil moisture availability relevant to drought-induced stress (Figs 1; S6). Thus, for the purpose of our analysis, we first identified the most severe drought events at each study site between 1970 and 2011, defined as those where CWD was 1.5 standard deviations below the annual site mean (Bose et al., 2024; Diffenbaugh et al., 2015; Tao et al., 2024). Of these, we then selected the most recent event in the timeseries, allowing us to directly link the impact of the drought to the plot census data collected in 2012. We also visually cross-checked the links between drought years with the detrended BAI time series for each study site to confirm that growth declines were associated with drought events rather than biotic disturbances such as bark beetle attacks or other pathogen outbreaks (Fig. 1; Schwarz et al., 2020). Based on this, we identified the following drought years at each site: 2002 in Poland; 2003 in Italy, Germany, and Romania; 2005 in Spain; and 2006 in Finland. These years correspond to well-known drought events that occurred across Europe at the start of the 21^st^ century, including the extreme 2003 heatwave that affected much of central Europe (Ciais et al., 2005) and the regional droughts that affected the Iberian Peninsula in 2005 (Jucker, Bouriaud, Avacaritei, Dănilă, et al., 2014) and Scandinavia in 2006 (Grossiord, Granier, Gessler, et al., 2014).

**Fig. 1:**
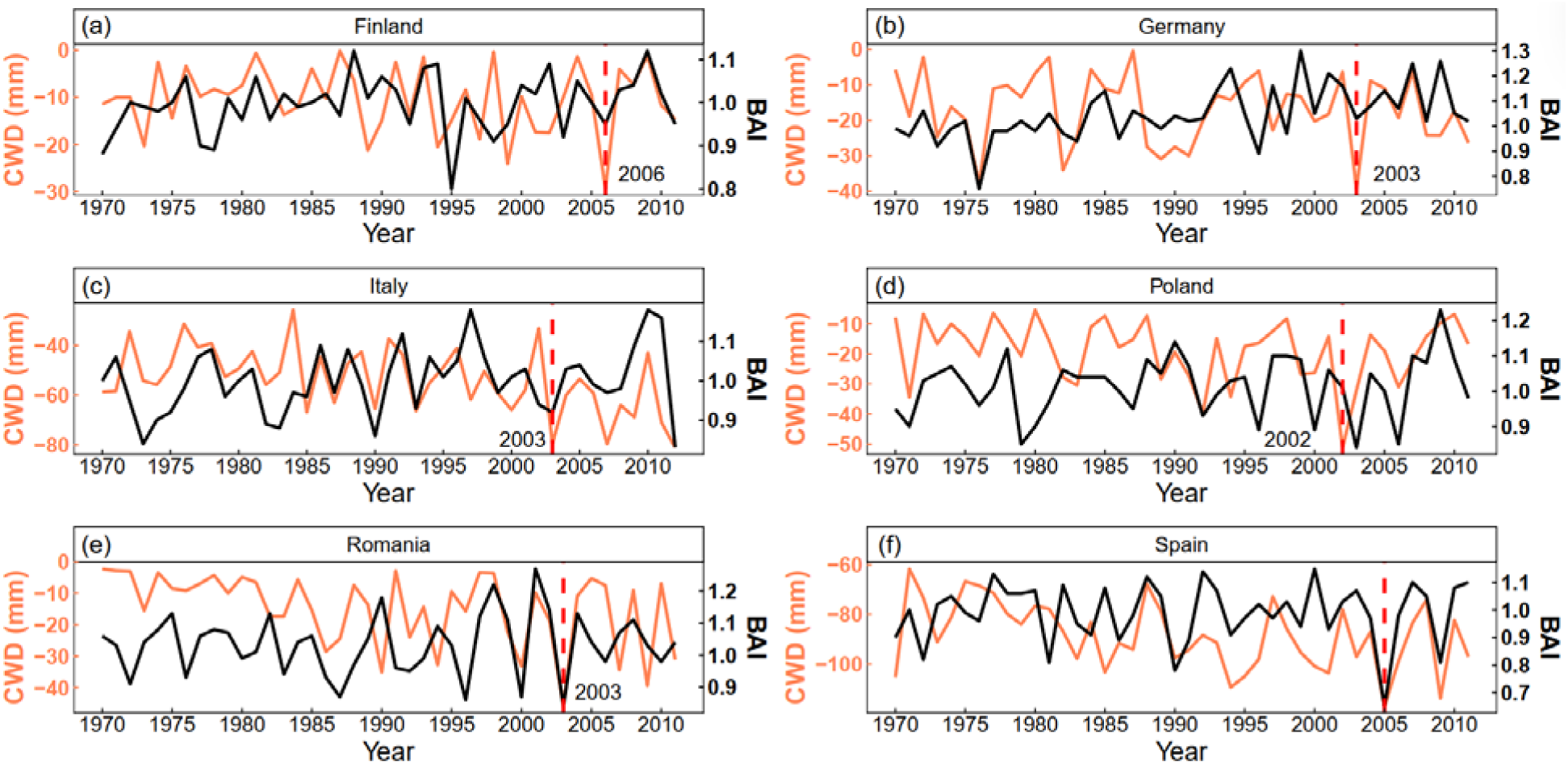
Identification of drought events. Drought years were identified separately for each FunDivEUROPE forest type between 1970-2011. This was done by identifying years with abnormally low growing-season climatic water deficit (CWD; in mm; orange lines) and comparing them with detrended basal area increment (BAI; unitless; black lines). When two or more candidate drought years were identified (e.g., 1976 and 2003 in Germany), the most recent drought was selected to better link it to the field census data collected in 2011-2012. The red vertical dashed line indicates the selected drought year for each forest type.

### Indices of tree growth responses to drought

Having identified drought years, we then used the tree ring data to calculate four widely used indices that capture how trees respond to and recover from drought (Lloret et al., 2011; Schwarz et al., 2020). These indices are computed by comparing a tree’s growth rate before (GRO_pre_), during (GRO_during_) and after drought (GRO_post_) and include resistance (Rt), recovery (Rc), resilience (Rs) and relative resilience (RRs):

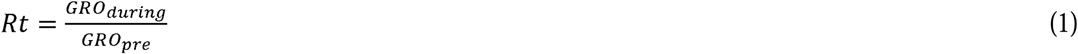

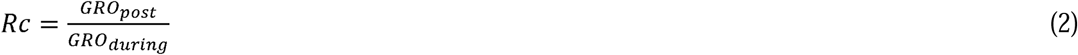

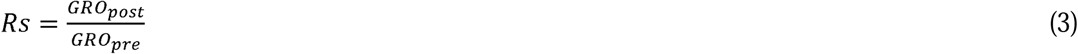

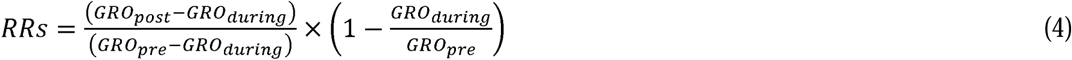

All indices were calculated using the BAI_detrend_ timeseries to avoid long-term ontogenetic growth trends confounding results (e.g., tendency of BAI to increase with tree size; Van Meerbeek et al., 2021). GRO_pre_ and GRO_post_ were defined as the mean BAI_detrend_ in the three years before and after the drought, respectively, as this is expected to encompass the legacy effects of drought in most species (Anderegg et al., 2015; Zang et al., 2014; Zhu et al., 2023), and because most trees recovered within two years (see Fig. S7). To better understand the relationship between these four indicators of tree responses to drought, we calculated pairwise Spearman’s rank correlations (ρ) between metrics (Fig. S8).

### Intrinsic and extrinsic predictors of tree responses to drought

We used both plot census and tree-ring data obtained between 2011 and 2012 to calculate a series of intrinsic and extrinsic factors known to influence tree growth and its response to drought. We chose to focus on these factors because they capture key features of a tree’s size and vigour, its competitive neighbourhood, and the drought event itself. Specifically, intrinsic factors included: (1) tree height (McGregor et al., 2021a), (2) tree age, calculated from the tree cores at breast height (Bigler et al., 2007; Marchand et al., 2025), (3) live crown ratio (LCR, defined as CD/H) as an indicator of crown dimension (Benavides et al., 2019), and (4) pre-drought growth rate, calculated as the mean growth in the 10 years before drought onset using the original, non-detrended BAI timeseries (Bose et al., 2020; Zhang et al., 2022). Extrinsic factors included: (5) stand-level tree species richness (SR), which ranged from 1–5 based on the FunDivEUROPE experimental design (Grossiord, Granier, Ratcliffe, et al., 2014; Jucker, Bouriaud, Avacaritei, & Coomes, 2014), (6) local competitive environment, quantified here as the cumulative basal area of trees taller than the target tree within 10 × 10 m subplots (stand basal area; Jucker et al., 2015; Vanderwel et al., 2020), and (7) drought intensity, defined as the CWD experienced by each plot during the growing season in the drought year (Hankin et al., 2019; Wu et al., 2022). Note that for a subset of trees where increment cores did not reach the pith (276 of 2909), tree age was estimated considering the first and last year of the ring-width series, using the *rwl.stats* function from *dplR* R-package.

### Modelling the drivers of tree-level resilience to drought

Prior to model fitting, we centred and scaled both predictors and response variables. We then used hierarchical Bayesian models implemented in the R package *brms* (Bürkner, 2017) to model variation in each of the four resilience indices (Rt, Rc, Rs, RRs). Predictor variables included the seven intrinsic and extrinsic factors described above as fixed effects, forest type (6 levels) nested within species (16 levels) as random intercepts with varying slopes (7 levels), and a random intercept term to account for additional unexplained variation among plots (208 levels). Due to the skewed distribution of the four response variables, we chose a skewed normal error distribution. Thus, for each resilience component, we fit the following model:

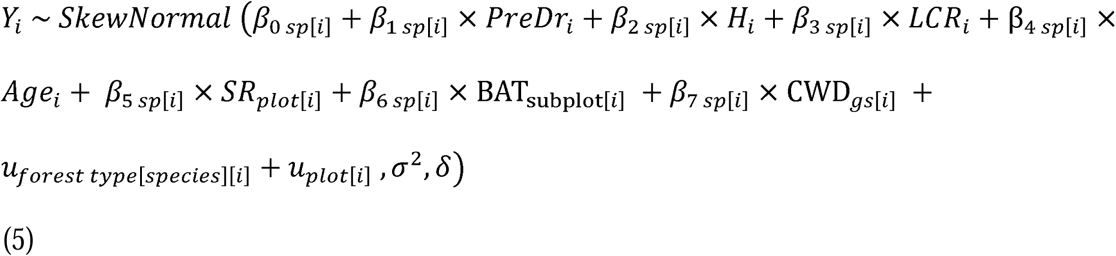

where Y is one of four resilience indices, *β*_0-7_ are fixed effects (intercept and slope), *sp*[*i*] refers to the fixed effects depending on the species identity of individual [*i*], *u*_forest type[species][i]_ is the random effect of forest type nested within species, and *u*_plot[i]_ is the random intercept term for the plot where individual [*i*] is located. The skewed normal distribution is parameterised through the scale parameter σ and shape parameter δ.

Hence, higher-level variation was modelled as follows:

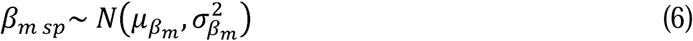

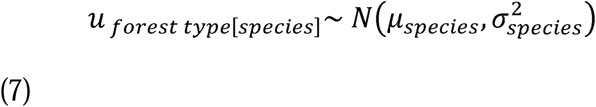

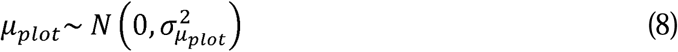

Lastly, to account for potential between-plot variation in the variance parameter, we explicitly modelled the scale parameter σ of the skewed normal distribution as a function of both species and plot as: *σ_i_*= *exp*(γ_0_ + γ*_plot_*_[*i*]_+ γ*_sp_*_[*i*]_) For all parameters and models, we chose weakly informative priors that constrained effects not to exceed the range of the response variables (i.e., normal distributions with *μ* = 0 and σ = 1.0 for fixed effects and *μ* = 1.0 and σ = 1.0 on log scales for the distributional parameters). Models were run using four independent chains, each with 2000 iterations (1000 warm-up iterations), with the default maximum tree depth of 10, and *adapt_delta* = 0.95. When models registered a small number of divergent transitions (<5) after warm-up, we increased both to 12 and 0.99, respectively. Model convergence was verified using the potential scale reduction factor (R-hat), with all values ≤ 1.01, and the bulk and tail effective sample sizes, with all values > 1000 (Tables S4-S7). Model fit was evaluated via posterior predictive checks, using the inbuilt *pp_check* function in *brms* (Fig. S2).

Following model fitting, we checked for multicollinearity among predictor variables by calculating variance inflation factors (VIFs) using the *check_collinearity* function in the *performance* package (Lüdecke et al., 2021) and confirming that VIFs were < 2 in all cases (Bose et al., 2020; Vanneste et al., 2024). We also used the *r2* function of the *performance* package to calculate the explained variance of each model, including both marginal (fixed effects only) and conditional *R*^2^ values (fixed and random effects combined) (Lüdecke et al., 2021). Standardised effect sizes were derived from posterior distributions of hierarchical Bayesian models. For the factor analysis (Fig. 2), posterior distributions of fixed-effect coefficients were extracted and summarised using posterior medians and two-sided 80% and 95% credible intervals (CIs) (Vanneste et al., 2024). Species-specific drought responses across forest types (Fig. 3) were quantified by extracting posterior distributions of varying intercepts, representing deviations from each species’ mean response. Figures, therefore, present posterior medians and associated uncertainty for both fixed and hierarchical (random) effects.

**Fig. 2:**
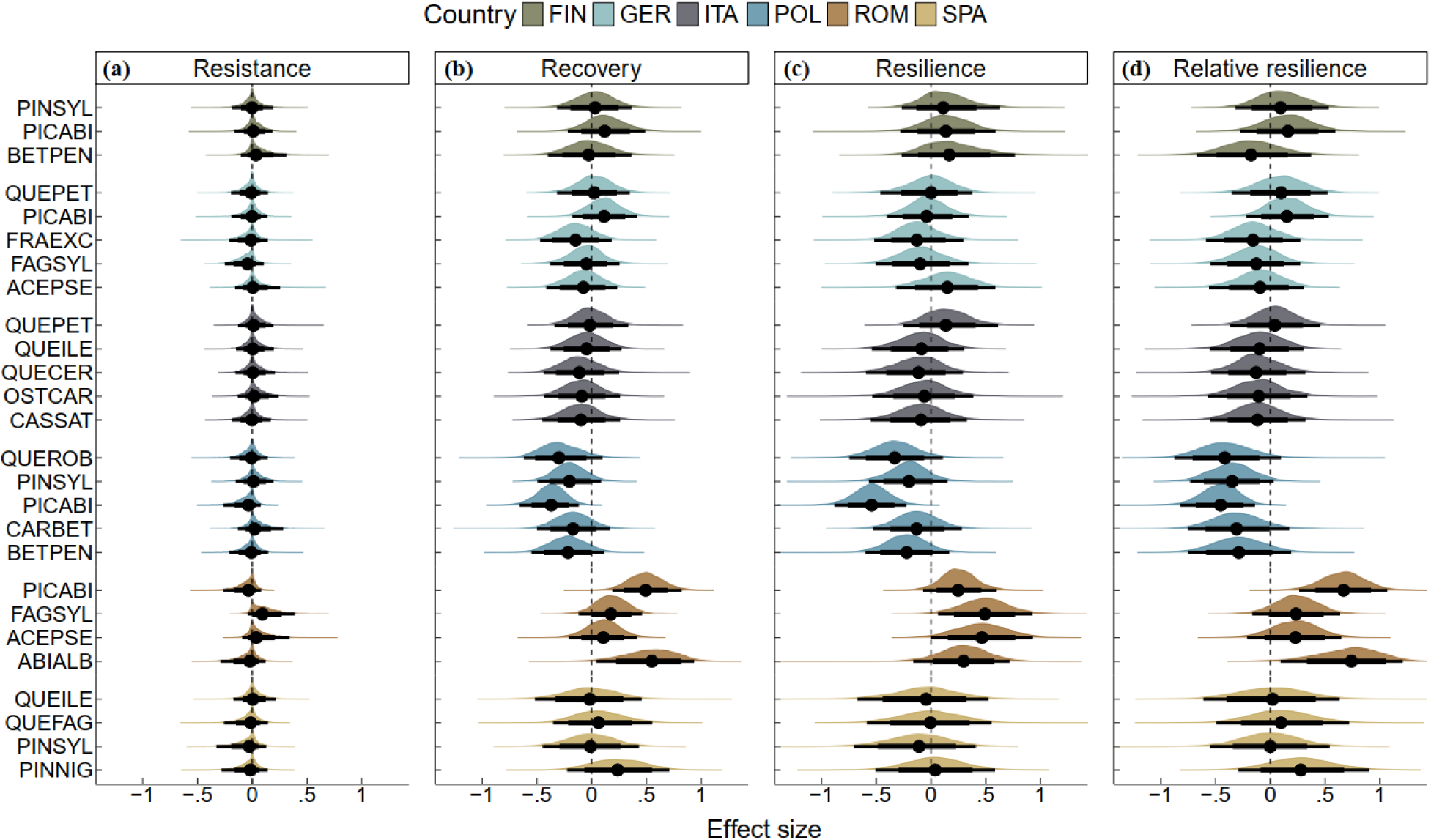
Variation in response to drought across tree species and forest types. Standardised effect sizes show posterior density median values (black circles) with 80% (thick bars) and 95% (thin bars) CIs for the effect of country nested within species on four components of overall tree growth resilience (resistance, recovery, resilience and relative resilience) of individual trees but when growing at different forest types (Finland, Germany, Italy, Poland, Romania and Spain). The dashed vertical line indicates the zero-effect size, and each density plot shows the posterior distributions of the parameter estimates obtained from the multivariate Bayesian model. The legend at the top shows the colours associated with each forest type (country) and the tree species growing within them. Species names are abbreviated using the first three letters of the genus and species name (see Table S3 for full species names).

**Fig. 3:**
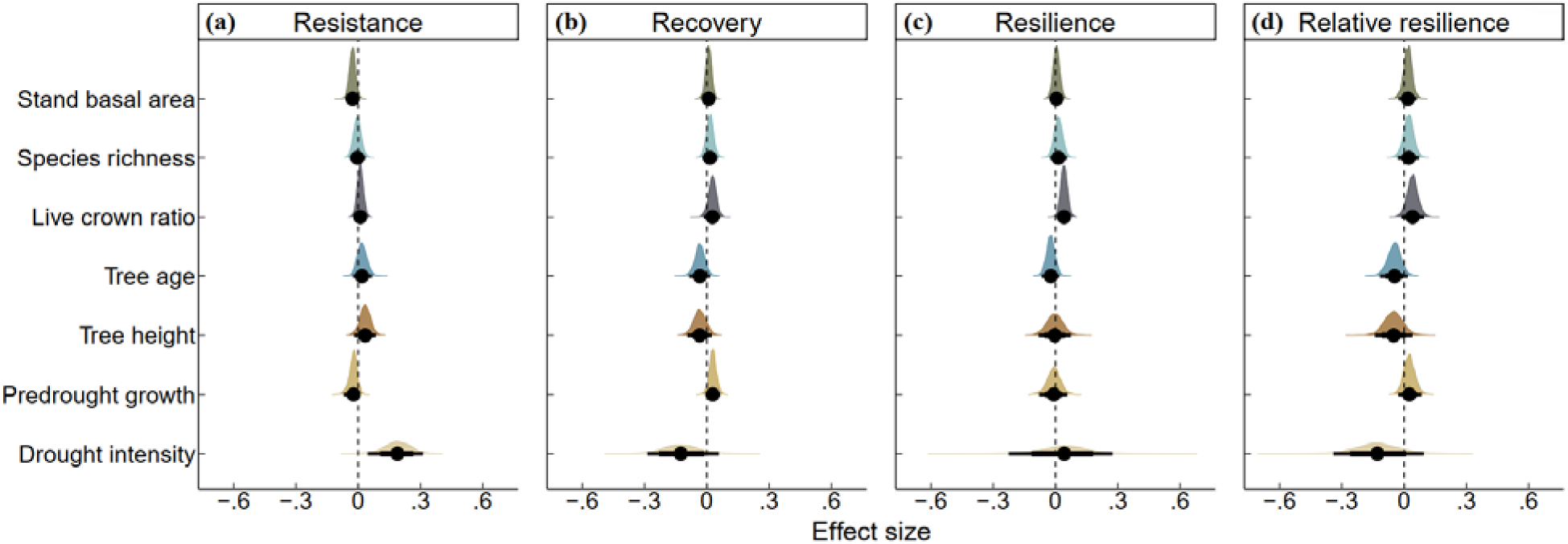
Influence of intrinsic and extrinsic factors on tree-level resilience to drought. Standardised effect sizes show posterior density median values (black circles) with 80% (thick bars) and 95% (thin bars) CIs for the effect of four intrinsic (predrought growth, height, live crown ratio and age) and three extrinsic factors (species richness, stand basal area and drought intensity) on four components of tree growth resilience (resistance, recovery, resilience and relative resilience) at the individual tree level. The dashed vertical line indicates the zero-effect size, and each density plot shows the posterior distributions of the parameter estimates obtained from the multivariate Bayesian model.

## RESULTS

### Variation in tree responses to drought among Europe’s major forest types and tree species

Overall, we found that tree species growing in Romania and Poland exhibited the highest and lowest resilience to drought, respectively (Fig. 2). Among gymnosperms, *Picea abies* and *Abies alba* consistently showed high recovery and relative resilience, whereas angiosperms such as *Fagus sylvatica* and *Acer pseudoplatanus* showed high resilience to drought. In contrast, most species from Poland exhibited the opposite trend, particularly *Picea abies*, which showed clear and consistent negative responses to drought (Fig. 2).

### Drivers of individual tree resilience to drought and its predictability

Across all resilience components, hierarchical Bayesian models explained a relatively low proportion of the variation in tree-level responses to drought (conditional *R*^2^ = 0.13–0.21; Table S8). Moreover, most of this variability was explained by differences among forest types, species identity, and plots (i.e., random effects), with the variability explained by the fixed-effect components of the models < 1% across all resilience metrics.

Of the intrinsic and extrinsic factors considered in our analysis, live crown ratio and drought intensity were the only two factors that had a clear effect on two resilience components (Figs 3-4). Specifically, trees with larger live crown ratios tended to exhibit greater drought resilience (mean [±□ 95% credible interval] □=□ 0.04; 0.01–0.07). While trees exposed to lower CWD during the growing season of the drought year exhibited higher resistance (0.19; 0.04–0.31). By contrast, we found no clear effects of pre-drought growth rate, tree age, or tree height on any component of growth resilience. Similarly, variation in a tree’s local competitive environment – defined in terms of both stand basal area and tree diversity – did not translate into clear differences in responses to drought (Fig. 3-4).

**Fig. 4:**
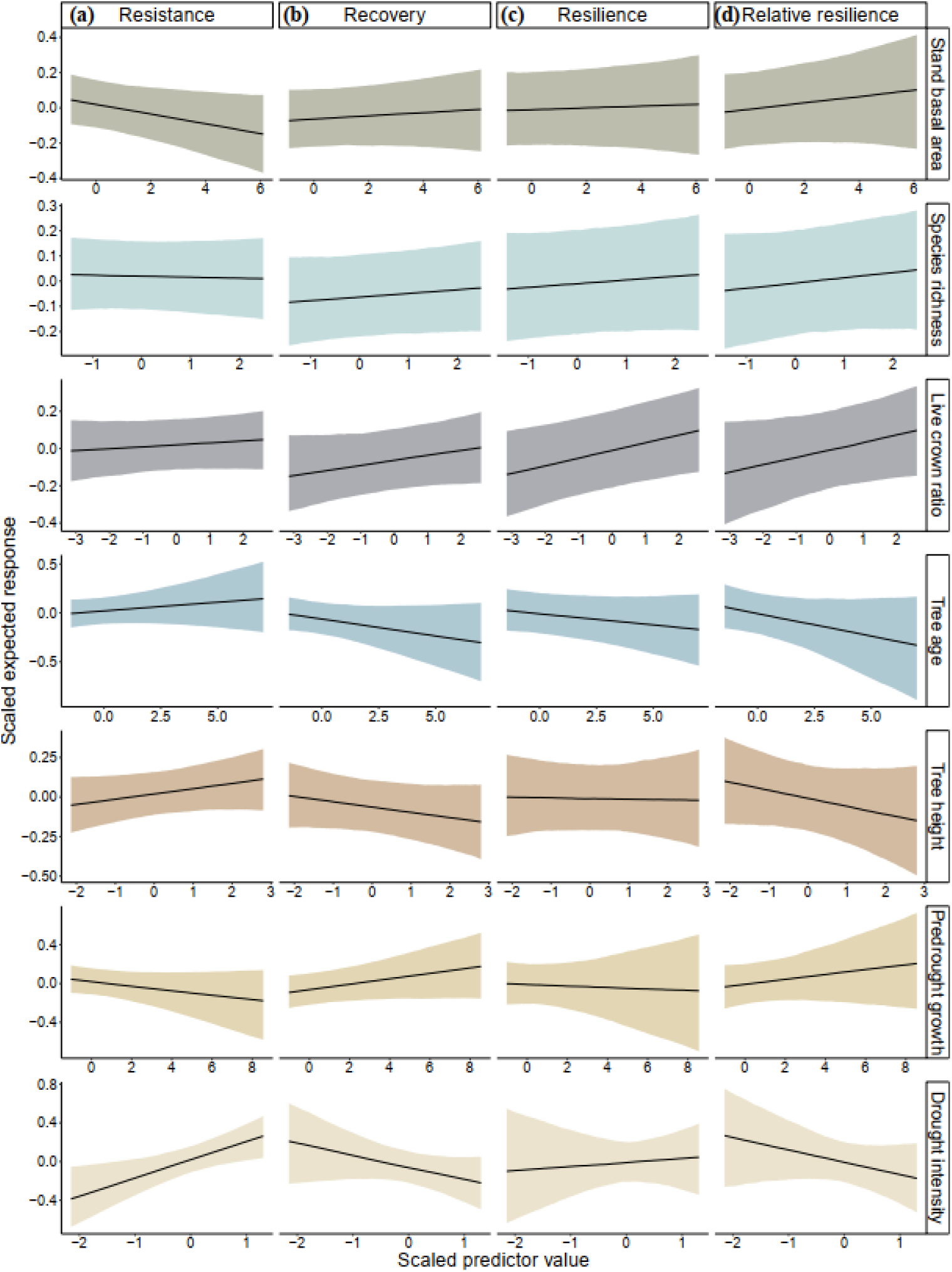
Expected response of tree resistance (Rt), recovery (Rc), resilience (Rs) and relative resilience (RRs) as a function of the seven predictors shown on the right axis of the panel. The black line represents the posterior mean estimate, with the shaded area indicating the 95% credible interval.

## DISCUSSION

### Limited predictability of individual tree responses to drought

Despite our comprehensive assessment of individual tree responses to drought, which included tree ring data from 2,909 individuals representing 16 major tree species distributed across six mature forest types in Europe, only live crown ratio emerged as a clear predictor of individual tree-level resilience. Even after incorporating drought intensity as another extrinsic predictor variable, the performance of our models remained low, explaining 13-21% of the total variance (Table S8). In Europe, previous investigations on the drivers of individual tree drought resilience have primarily focused on the most common and productive tree species and are often restricted to a single provenance or region (Bottero et al., 2021; Gazol & Camarero, 2016; Martínez-Vilalta et al., 2012; Pretzsch et al., 2013; Zang et al., 2014). Despite differences in species coverage and modelling approach, such as the tendency of many studies to fit more complex models and to fit them separately for each species due to limited species representation, these studies consistently reported low explanatory power in predicting tree-level resilience, aligning with our findings. This underscores the inherent difficulty of predicting drought resilience at the individual-tree level, stressing the need to consider the cumulative drought history of trees rather than focusing on single major events when modelling their drought responses (Camarero et al., 2024; Song, Sterck, Sass Klaassen, et al., 2022; Sterck et al., 2024). It also highlights the importance of developing integrative tools that leverage tree-ring, physiological, and remote-sensing data with mechanistic forest simulation models to better capture non-linear and delayed growth responses to drought stress.

### No effect of species richness or competition on tree responses to drought

Overall, we found that a tree’s local competitive environment – characterised both in terms of its density and diversity – had little discernible impact on its response to drought. In particular, species richness did not confer greater drought resilience in our global models, showing no consistent effects across species (Forrester et al., 2016; Grossiord, Granier, Ratcliffe, et al., 2014) or among individual trees (Figs 3-4; Merlin et al., 2015; Sachsenmaier et al., 2024; Serrano León et al., 2025). This is likely due to the limited tree diversity gradient within our study plots, as across the entire FunDivEUROPE forest plot network, mixed stands typically contain no more than four species, with only a single plot in Poland reaching a maximum of five within a 30 × 30 m area. Previous studies that reported significant effects of species richness or functional diversity on drought resilience—such as those of Martínez-Vilalta et al. (2012), Gazol & Camarero (2016), and Fichtner et al. (2020) —were conducted under more constrained conditions. The first two focused on two single well-known species—*Pinus sylvestris* and *Abies alba* —while the latter employed an experimental setup with 40 sapling species in a subtropical Chinese forest, limiting comparability with mature, mixed-species temperate forests.

Martínez-Vilalta et al. (2012) found a significant effect on BAI in *Pinus sylvestris* when treating species richness as a discrete variable with three levels: 1, 2–3, and 4–8 species; and indeed, the only clear effect size was observed at the third level, where there were more species. In line with this, we also found a clear positive effect of species richness on recovery and relative resilience, and a negative effect on resistance of *Pinus sylvestris* from the same origin, as well as a positive effect on the resilience of *Quercus robur* trees from Poland (Tables S23-S26), suggesting that the effects of species richness are species-specific (see also Jucker et al., 2015). Similarly, Gazol & Camarero (2016) did not find significant effects of inter- or intraspecific competition, and reported only a weak positive relationship between functional diversity and the growth resilience of *Abies alba* to the 1985–86 Spanish drought.

Additionally, Fichtner et al. (2020) only found a negative effect of neighbourhood competition and a weak positive effect of species richness. However, this research was conducted as a controlled experiment with a higher species density (N= 40) and at the sapling level (6 years old). In addition, recent work shows that although species richness alone seldom enhances drought resilience, functional diversity can acquire predictive power when interacting with drought duration, with stronger neighbourhood effects emerging under prolonged dry periods (Serrano León et al., 2025). Thus, the influence of competition and species richness remains to be further explored using a wider set of tree species and forest types. Furthermore, it’s also important to take into consideration other facets of diversity (e.g., phylogenetic), as recent evidence from subtropical China suggests that these components of diversity can have a stronger impact on ecosystem functioning and temporal stability (Liang et al., 2024; Rodríguez-Hernández et al., 2021).

### Live crown ratio mitigates the impact of drought-induced stress in trees

Although some studies have evaluated the influence of crown conditions on tree growth during (Camarero et al., 2015) and after (Galiano et al., 2012; Schmied, Pretzsch, et al., 2023) a given drought episode, fewer studies have ever tested the influence of crown dimensions (e.g., width or depth) on individual tree-level responses to drought. In our research, we consistently observed a small positive effect size of LCR on Lloret’s resilience components, except for resistance, where it was more costly for species to retain a larger number of leaves relative to their height (Fig. 4a). Accordingly, we observed the strongest recovery, resilience, and relative resilience in trees with LCR >30 (Figs 3; S3a; Stăncioiu et al., 2021), though this effect is more evident in *Carpinus betulus* and *Picea abies*, which have larger LCRs than the other species (Fig. S3a). This suggests that the small overall effect size of LCR may be attributed to its high variability across and within species, regardless of age (Fig. S3a,b). Further, LCR does not account for width or spread, and length does not always accurately reflect crown volume (Stăncioiu et al., 2021). Hence, a variable that combines both crown width and depth (e.g., between crown shape and live crown ratio), as well as the shadow generated by the crowns of taller individuals (Ma et al., 2023), before, during, and after a given drought episode, could help better understand crown architecture’s influence on tree resilience to drought.

### Drought resiliency is more strongly driven by forest type than by species identity

Unlike Bose, Gessler, et al. (2020), we found tree resilience to drought also depended on the geographical location where trees were growing (Figs 2; S4; Gazol et al., 2017; Zhu et al., 2023). This implies that species growing at different geographical locations (e.g., longitude, latitude and altitude) and forest types respond differently to climate extremes (Figs 2; S7-S8; Bose et al., 2020; Carnwath & Nelson, 2017; Zhu et al., 2023). Contrary to our expectations, we found that tree responses to drought depended more on forest type than on species identity (Figs 2; S5; Serra-Maluquer et al., 2018). Generally, we found that trees growing in Romania and Poland exhibited the highest and lowest levels of resilience, respectively. In contrast, trees from Spain were the least resistant, likely due to their highly limited water availability, which is even more pronounced during summer spells (Fig. S6). Poland, home to Europe’s last virgin forest (Białowieża), surprisingly suffered the most from drought, signalling that this forest ecosystem is more threatened if extreme droughts persist in the short term. A possible explanation for this low resilience is that Poland has experienced a temperature increment of 1.27 °C from 1950 to 2015, leading to a higher frequency of droughts, which in turn has promoted bark beetles and caused massive diebacks on *Picea abies* individuals (Boczoń et al., 2018), the most dominant tree species growing in this area. Indeed, *Picea abies* showed the highest drought sensitivity, though all Polish species in our data set exhibited similar negative responses to drought (Figs 2; S4).

Additionally, we found contrasting species-specific responses to drought across the six forest types studied, with the clearest patterns found in *Picea abies*, *Abies alba* and *Fagus sylvatica* (Fig. 2). While these differences highlight some degree of species-level sensitivity, these were limited to only these three species out of a total of sixteen present in our data set, which suggests that the unique topographic features and climatic conditions of each type of forest play an even greater role in shaping tree growth and their responses to drought at the level of individual trees (Bose et al., 2020; Carnwath & Nelson, 2017). Accordingly, trees growing in Romania, often on consistently steep slopes, tended to show higher drought resilience (Gazol et al., 2017), whereas *Picea abies* and other species from Poland, typically growing in lowland terrain, appeared more drought-sensitive (Figs 2; S4, S7). This pattern aligns with recent findings at the tree level by Thomas et al. (2024), who reported lower mortality risk for trees growing on upper hilly slopes compared to forests on flatter terrains. Our findings therefore suggest that common European species such as *Picea abies* and *Fagus sylvatica* may be more vulnerable in certain forest types and face increased replacement pressure from more drought-tolerant species, consistent with previous projections (Lévesque et al., 2013).

### Pre-drought growth rate does not affect how trees respond to drought

Unlike previous studies (Bose et al., 2020; Martínez-Vilalta et al., 2012; Zang et al., 2014; Zhang et al., 2022; Zhu et al., 2023), we found no clear effect of pre-drought growth rate across any component of tree resilience (Figs 3-4). A possible explanation for this effect is that trees can exhibit either fast or slow average growth rates preceding drought onset, which makes the effect of pre-drought growth less important when a broader range of species and individuals is considered. Accordingly, we found most species showed signs of a strong recovery capacity, as recovery was generally unaffected by pre-drought growth rate once conditions returned to normal (Martínez-Vilalta et al., 2012; Schmied, Pretzsch, et al., 2023; Tao et al., 2024), consistent with the average recovery period, which took no more than 1.5 years following drought impact (Fig. S7). Another stronger reason is that here we did not include detrended pre-drought growth as a predictor in our models, as most previous studies have done (Bose et al., 2020; R. Liang et al., 2024; Zhang et al., 2022), because doing so would not capture the actual growth variability among individual trees prior to drought.

### Tree height and age do not influence the drought resilience of individual trees

Overall, we found tree height and age did not influence any component of tree resilience (Figs 3-4; Tables S4-S7). This suggests that the effect of tree height on drought resilience is not necessarily controversial, but rather difficult to disentangle at the individual level, and thus probably more reliant on other species-specific functional traits (Fernández de Uña et al., 2023; González de Andrés et al., 2021). Similarly, although age has been shown to significantly influence the drought resilience of individual trees, due to growth rates tend to decline as trees age (Zang et al., 2014). Here, we could not find any significant effect of age on any component of resilience, suggesting that neither tree age nor height is a strong driver of tree-level resilience (Zhang et al., 2022; Zhu et al., 2023). Additionally, when we analysed the effects of height on each tree species individually, we found that the impact was noticeable only in a few fast-growing species across various resilience metrics. In most cases, the effect was negative, except for resistance. This influence was observed in six out of the sixteen species in our data set, including: *Betula pendula* from Finland, *Carpinus betulus* and *Picea abies* from Poland, *Fagus sylvatica* and *Fraxinus excelsior* from Germany, *Abies alba* and *Picea abies* from Romania (Tables S9-S22). We therefore believe the effect of height may not be detectable for several reasons: 1) here we show that predicting drought resilience at the tree-level is challenging, 2) we show that height affects only certain species irrespective of age, as we have trees covering all age/size classes throughout the entire chronology, 3) tree height is just a single dimension that may interact with root, stem, and crown dimensions, 4) we did not categorize height into different age/size classes as previous studies have done to overpredict resilience (Au et al., 2022), 5) we only measured height during one time period, meaning we lack real height measurements from historic drought events where we observed trees tended to suffer more from drought, and 6) we did not compare its effects across mild and severe drought episodes (Sachsenmaier et al., 2024). We conclude that the effects of tree height and age may be more apparent at the species level (Bennett et al., 2015; McGregor et al., 2021; Stovall et al., 2019) rather than at the tree level and thus warrant further investigation using more species-specific functional trait data.

### Drought intensity influences the ability of trees to maintain growth, but not their recovery from drought

In line with work by Zang et al. (2014) and more recent research by Zhang et al. (2022) conducted at the individual tree level, we found consistent patterns in how drought intensity influenced the different components of tree resilience (Figs. 2-4). Along the growing season, tree resistance increased with decreasing drought intensity (i.e., less negative CWD values). By contrast, tree recovery and relative resilience were higher under more intense drought stress (Figs. 3-4). For overall resilience, however, we found no consistent patterns, as the mean effect size was close to zero, suggesting that trees can exhibit both positive and negative growth responses during intense drought (Zang et al., 2014).

A possible explanation for the unclear effect of drought intensity on the other components of resilience is that individuals of different tree species exhibit varying growth strategies in response to drought stress—for example, the clear trade-off between conifers and broadleaved species (DeSoto et al., 2020). In addition, Gazol et al. (2017) found that as drought intensity (measured as the difference between soil moisture and potential evapotranspiration) increases, tree resistance declines, while post-drought recovery improves. This opposing pattern weakens the overall impact of drought intensity on tree resilience observed here and in previous studies (Gazol et al., 2017; Zhang et al., 2022). It is important to note that, as in most previous studies, drought stress—here quantified as the growing-season climatic water deficit (CWD) during the drought year—was measured at the plot level. This approach may overlook individual-tree responses, as they experience water stress differently depending on their specific microsite conditions.

## CONCLUSIONS

In a comprehensive latitudinal study spanning a wide range of environmental conditions, six mature natural forest types, and sixteen major tree species from Europe, we found limited predictability of intrinsic and extrinsic factors upon individual tree-level resilience to drought, as our models could only explain up to 21% of the total variance across the four resilience components. We found that trees with larger living crowns were generally better able to recover from drought and showed higher resilience, while trees exposed to lower drought intensities were better able to withstand drought. Furthermore, drought resilience did not depend on species identity but rather on the forest type and specific topographic conditions in which trees were growing. Despite our comprehensive assessment of tree responses to drought across Europe, we found only live crown ratio and drought intensity had clear evidence as predictors of tree-level resilience. This highlights the need to consider the cumulative drought history trees have experienced throughout their lifecycles when modelling their drought responses. Additionally, it pinpoints the need to develop next-generation drought-monitoring tools that integrate tree-ring, physiological, and remote-sensing data with mechanistic models, together with more robust, multi-event-based resilience metrics capable of capturing long-term stress accumulation to safeguard European forests against the impacts of climate change.

## Supporting information

ESM_Drivers of tree resilience

## ACKNOWLEDGEMENTS

The FunDivEUROPE project was funded through the European Union Seventh Framework Programme (grant: 265171). DIRH was funded through a Scholarship from the Consejo Nacional de Ciencia y Tecnología (CONACyT) of Mexico (grant: 809844). TJ was supported by a UK NERC Independent Research Fellowship (grant: NE/S01537X/1) and by a UKRI Frontier Research award (grant: EP/Y003810/1).

## DATA AVAILABILITY STATEMENT

All data and R code underpinning the results of this study will be publicly archived on Zenodo following the review of this paper.

