## Supplementary material for "Limited predictability of tree-level responses to drought across European forests": ESM_Drivers of tree resilience

Electronic supplemental material for:

**FEM**

By: Diego I. Rodríguez-Hernández^1*^, Fabian J. Fischer^1^, Duncan A. O’brien^1^, Martin De Kauwe^1^, Bo Wang^2^, Olivier Bouriaud^3^ and Tommaso Jucker^1^

^1^*School of Biological Sciences, University of Bristol, 24 Tyndall Avenue, Bristol, BS8 1TQ, UK*

*^2^State Key Laboratory of Cryospheric Science, Northwest Institute of Eco-Environment and Resources, Chinese Academy of Sciences, Lanzhou, 730000, China*

^3^*Stefan cel Mare University of Suceava, Str. Universitătii 13, 720229 Suceava, Romania*

**Figure S1.** Examples of Friedman’s supersmoother spline detrending method applied to BAI.

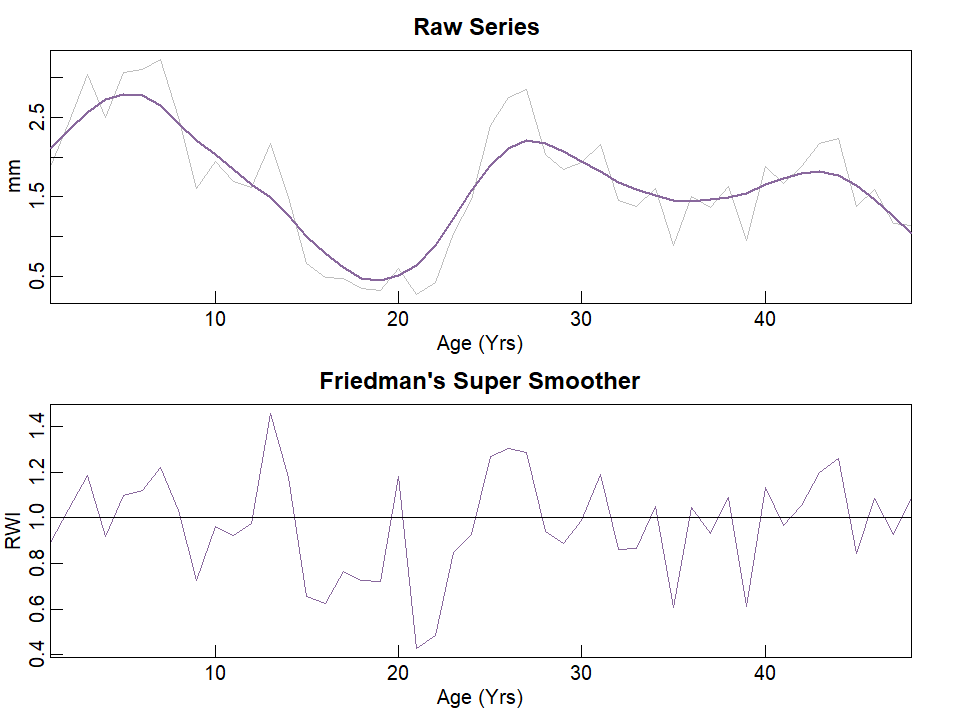

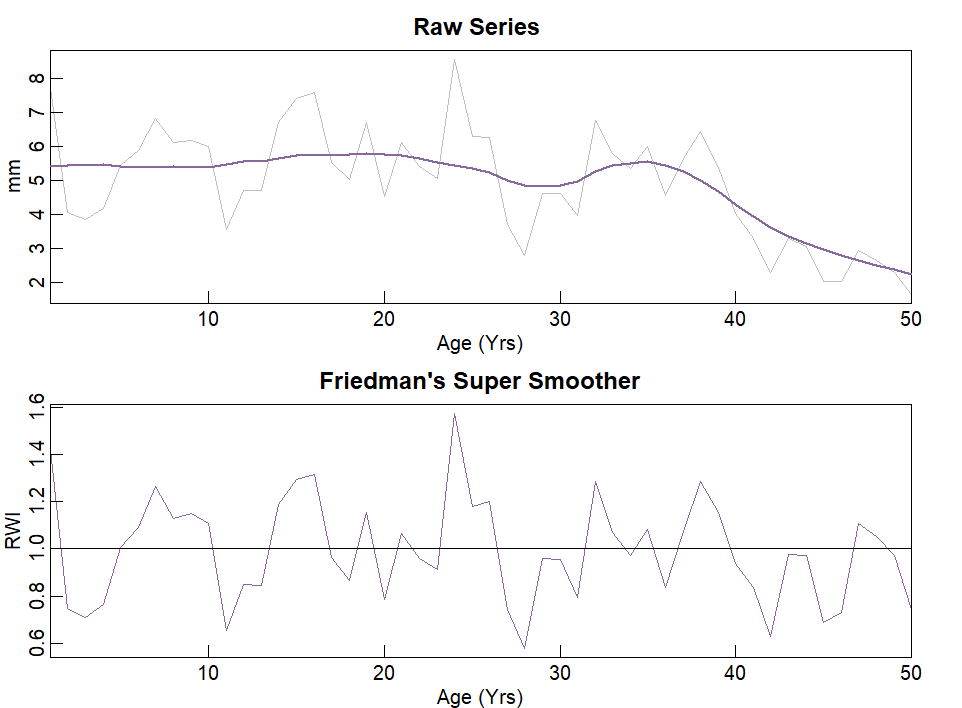

**Figure S2.** Posterior predictive checks (*pp_checks*) for the hierarchical Bayesian mixed models testing the effect of four intrinsic (pre-drought growth, height, live crown ratio, age) and three extrinsic variables (species richness, stand basal area, drought intensity), country (6 levels) nested within species (16 levels) and plot (208 levels) as random effects on four resilience indices (resistance, recovery, resilience, relative resilience). Based on the posterior distribution of the model parameters, 100 posterior draws (**y_rep_**) were simulated for each growth resilience index, and their densities were plotted against the density of the observed data (**y**). Model fits are considered accurate when the light blue lines (**y_rep_**) overlap with the dark blue line (**y**).

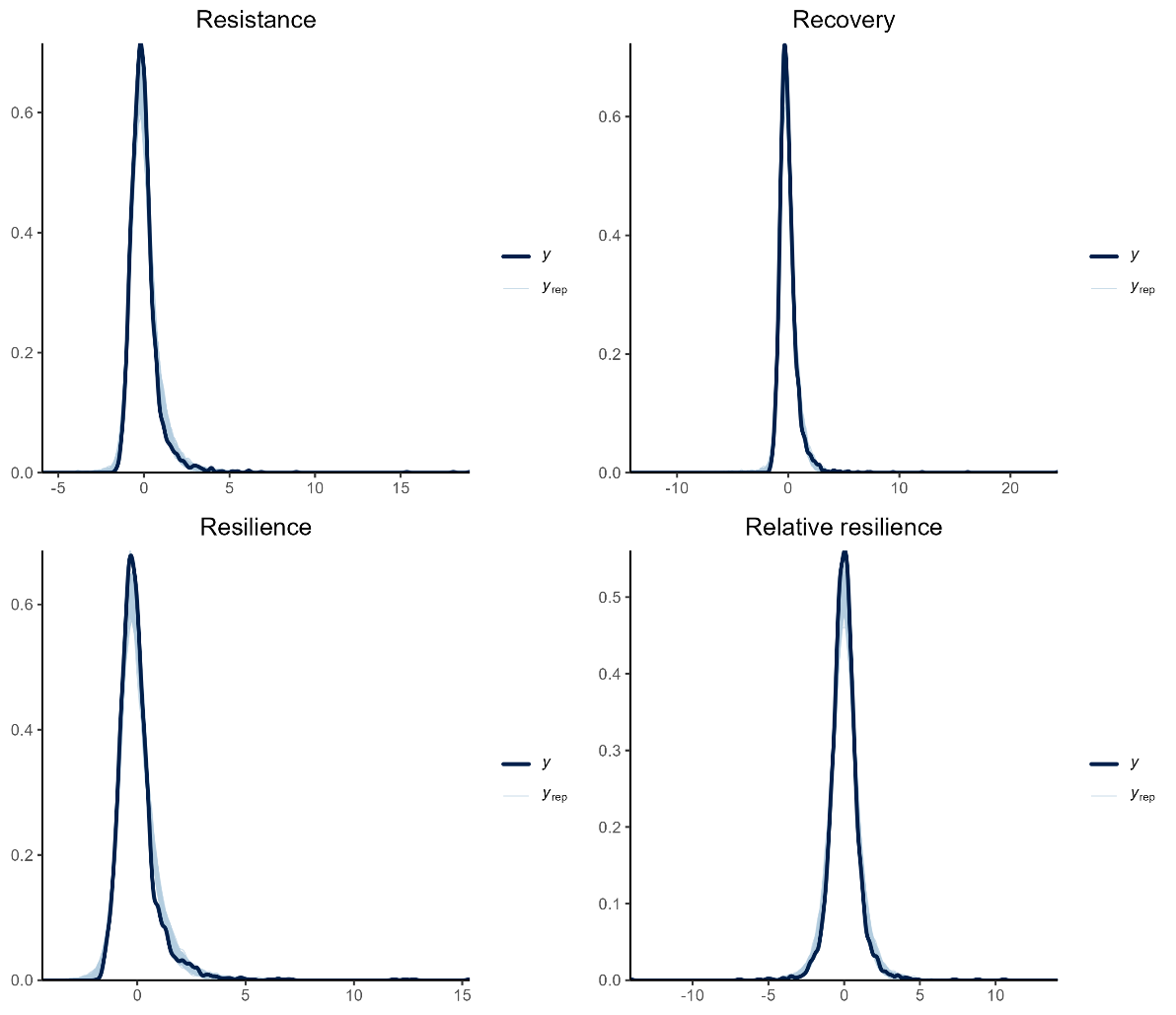

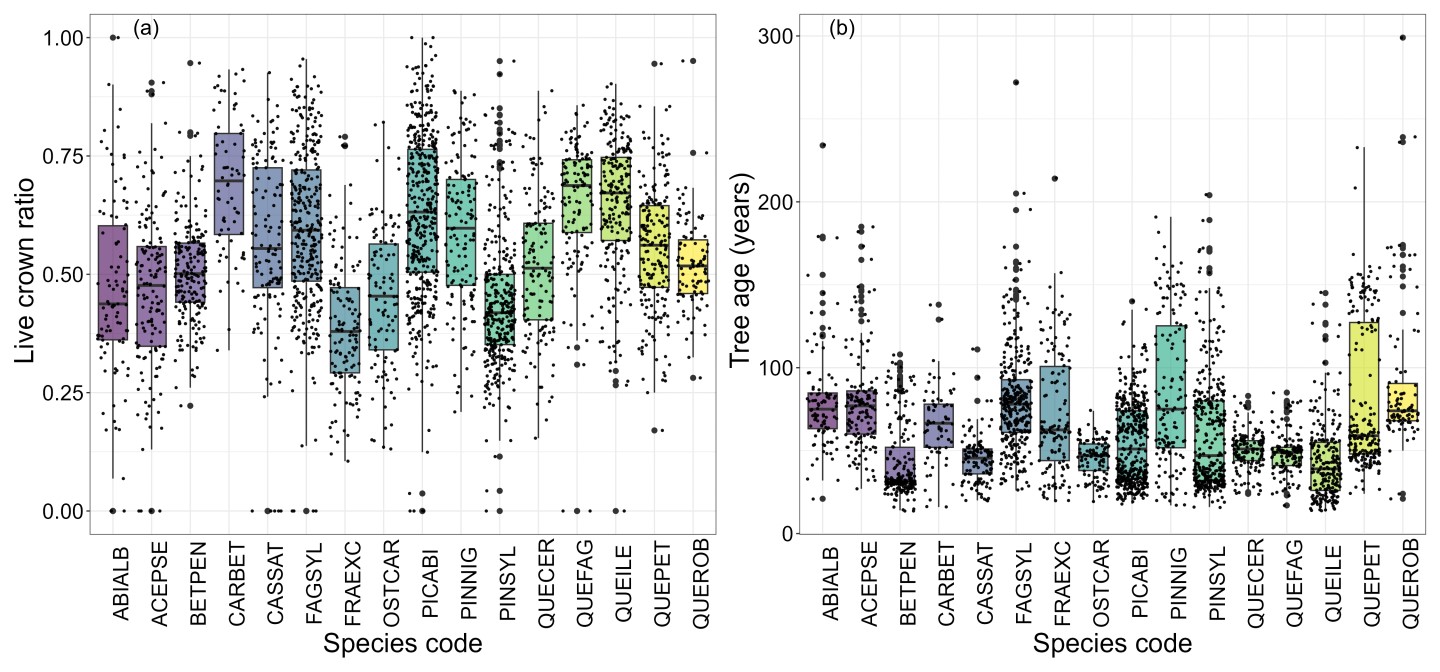
**Figure S3.** Box plots depicting the actual distributions of live crown ratio (unitless) and tree age of all individuals within each tree species from the FunDivEUROPE forest plot network. Box limits indicate the range of the central 50% of the data, with a central line marking the median value. Lines extending from each box capture the range of the remaining data, with bigger dots placed past the line edges to indicate outliers. The x-axis shows species code, while the y-axis the corresponding values for each structural attribute. For full species names, we refer the reader to Table S3.

**Figure S4.** Country-level resilience in response to drought. Standardised effect sizes show posterior median values (black circles) with 80% (thick bars) and 95% (thin bars) CIs for the effect of country as a random intercept term (Finland, Germany, Italy, Poland, Romania and Spain) on five resilience components (resistance, recovery, resilience, stability and lag-one autocorrelation) of individual trees belonging to 16 species and 26 populations. The dashed vertical line indicates the zero-effect size, and each density plot shows the posterior distributions of the parameter estimates from the multivariate Bayesian model.

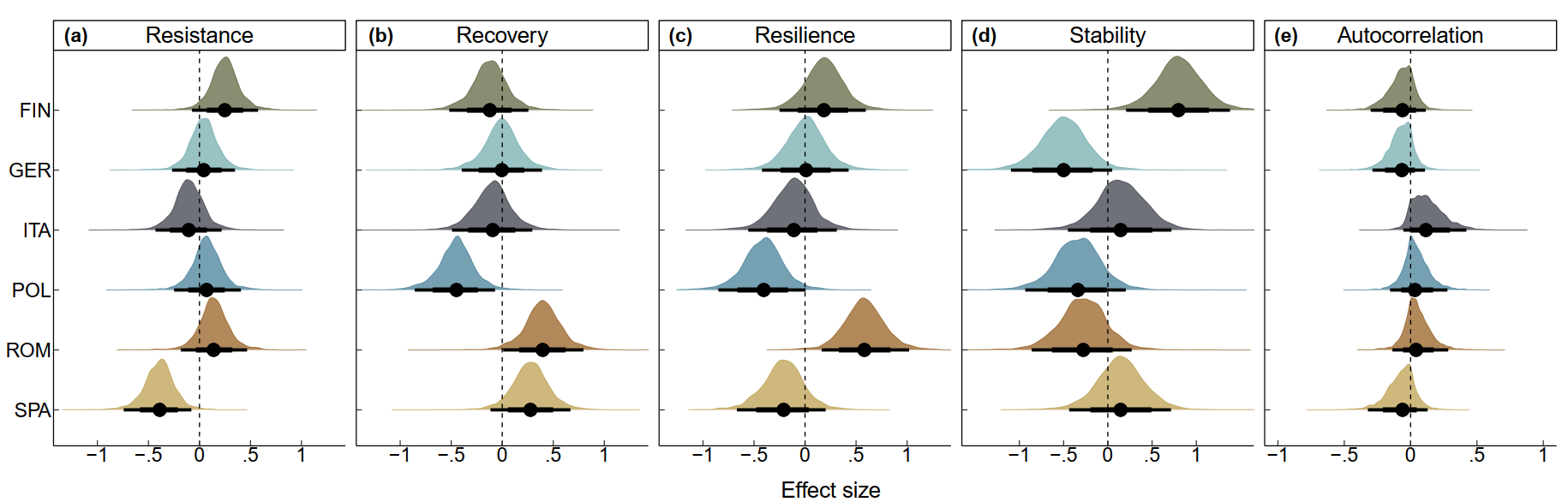

**Figure S5.** Species-identity responses to drought. Standardised effect sizes on the x-axis show posterior median values (black circles) with 80% (thick bars) and 95% (thin bars) CIs for the effect of four intrinsic (pre-drought growth, height, live crown ratio, age) and three extrinsic variables (species richness, stand basal area, drought intensity) on four resilience indices [from left to right: resistance (Rt), recovery (Rc), resilience (Rs), relative resilience (RRs)] of individual trees. The dashed vertical line indicates the zero-effect size, and each density plot shows the posterior distribution of the parameter estimates from a multivariate Bayesian model. For full species names, we refer the reader to Table S3.

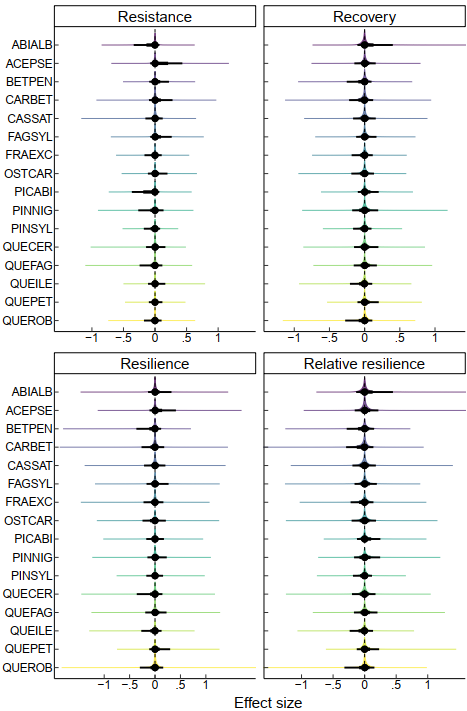

**Figure S6.** Annual seasonality of climatic water deficit (CWD; mm) for each forest type of the FunDivEUROPE forest plot network (FIN=Finland, GER=Germany, ITA=Italy, POL=Poland, ROM=Romania, SPA=Spain) between 1970 and 2011. The legend on the right-hand side shows the CWD across the different seasons of the year (spring, summer, autumn, winter and growing) represented by different viridis line colours
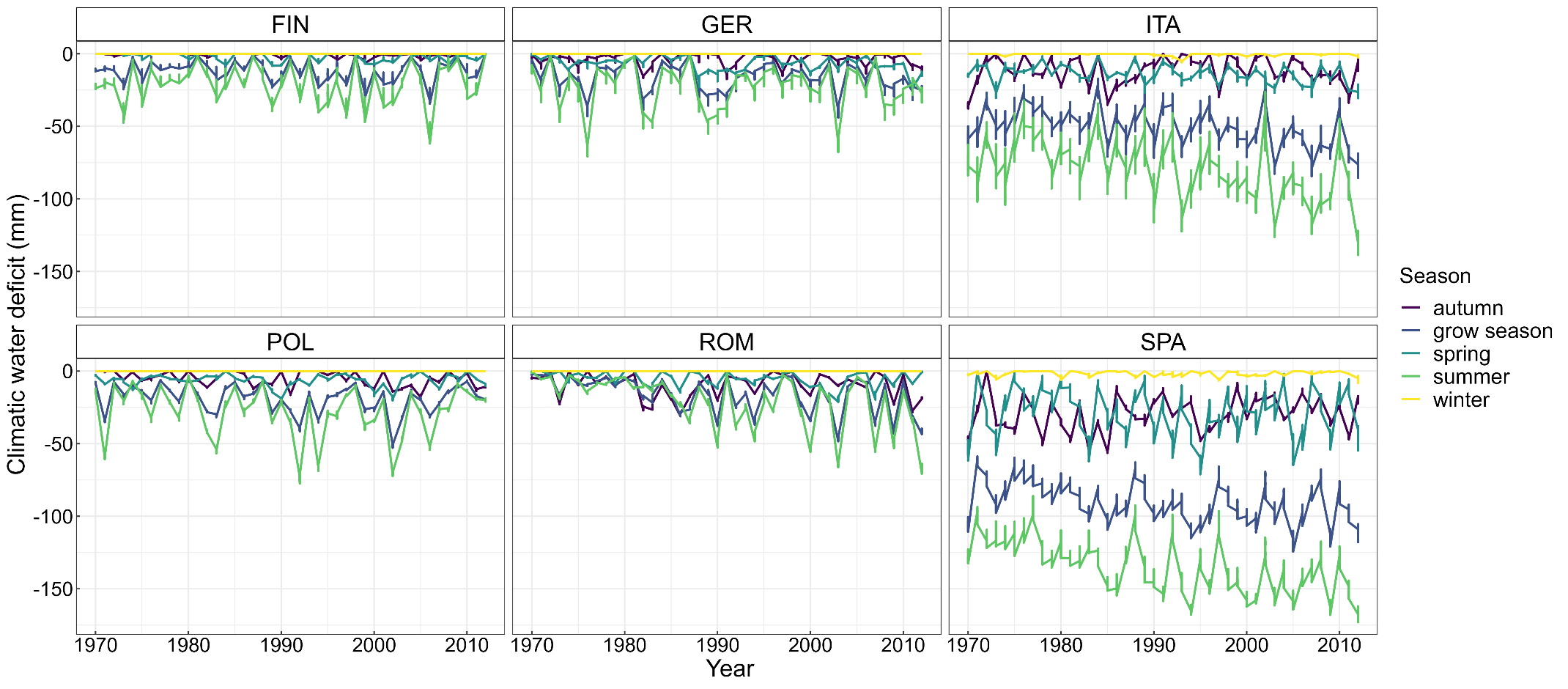
.

**Figure S7.** Descriptive statistics (mean ± SE) in response to drought for the 26 study populations (16 species) from the FunDivEUROPE forest plot network. The x-axis displays the mean values (circles) and standard error (bars) for each species’ resilience component, while the y-axis species code. The average recovery period (RcP) is expressed as the number of years needed to recover from drought impact, with the dotted vertical line representing the average RcP across the entire community. Different colours represent species growing in different countries, as shown in the legend. For full species names, we refer the reader to Table S3.

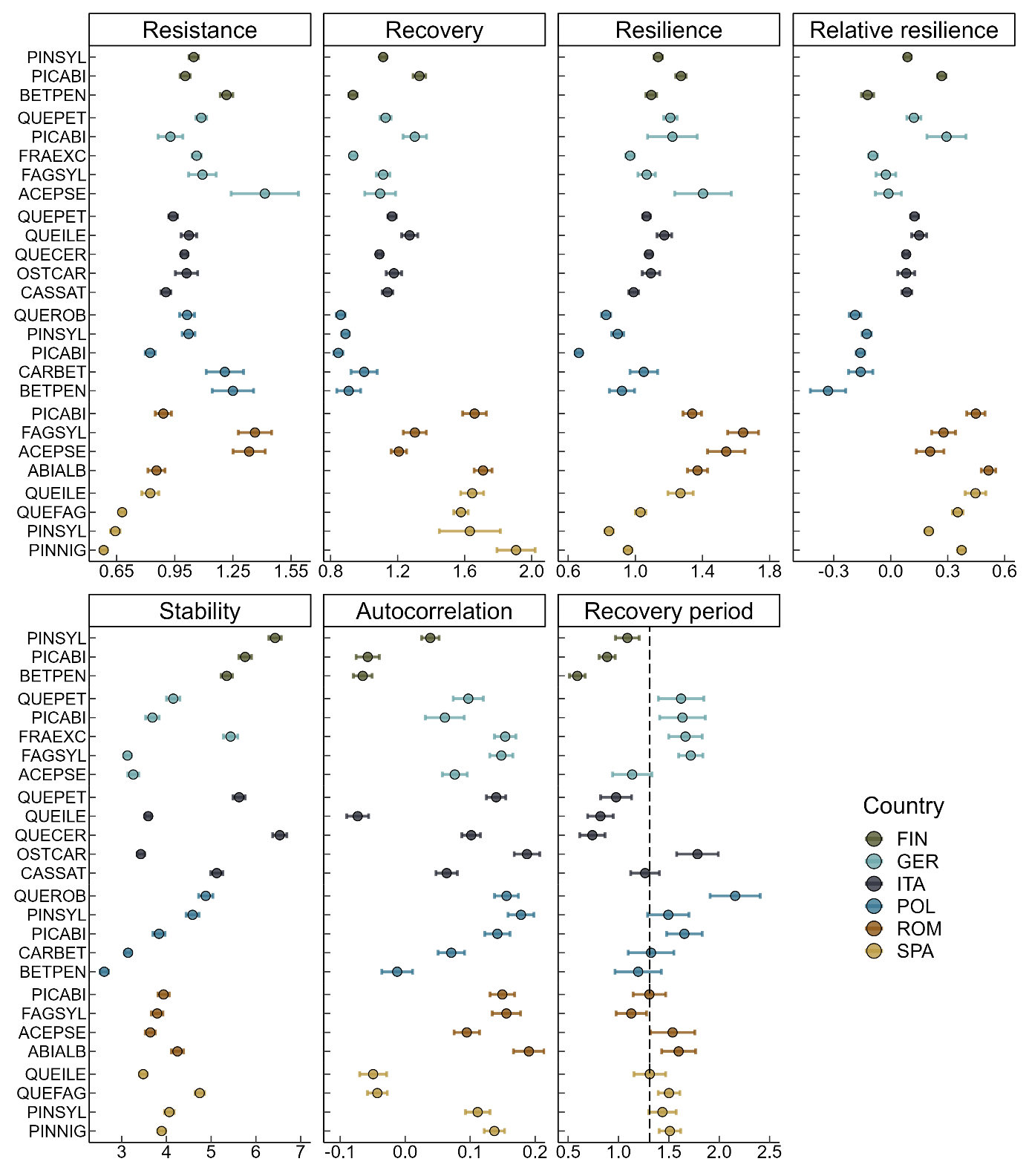

**Figure S8**. Spearman’s rank pairwise correlation (ρ) matrix among four indicators of tree responses to drought. Fitted lines correspond to a LOESS spline, while the density plots along the diagonal show the distribution of each metric. Note that resistance, recovery, and resilience were log-transformed due to their left-skewed distributions.

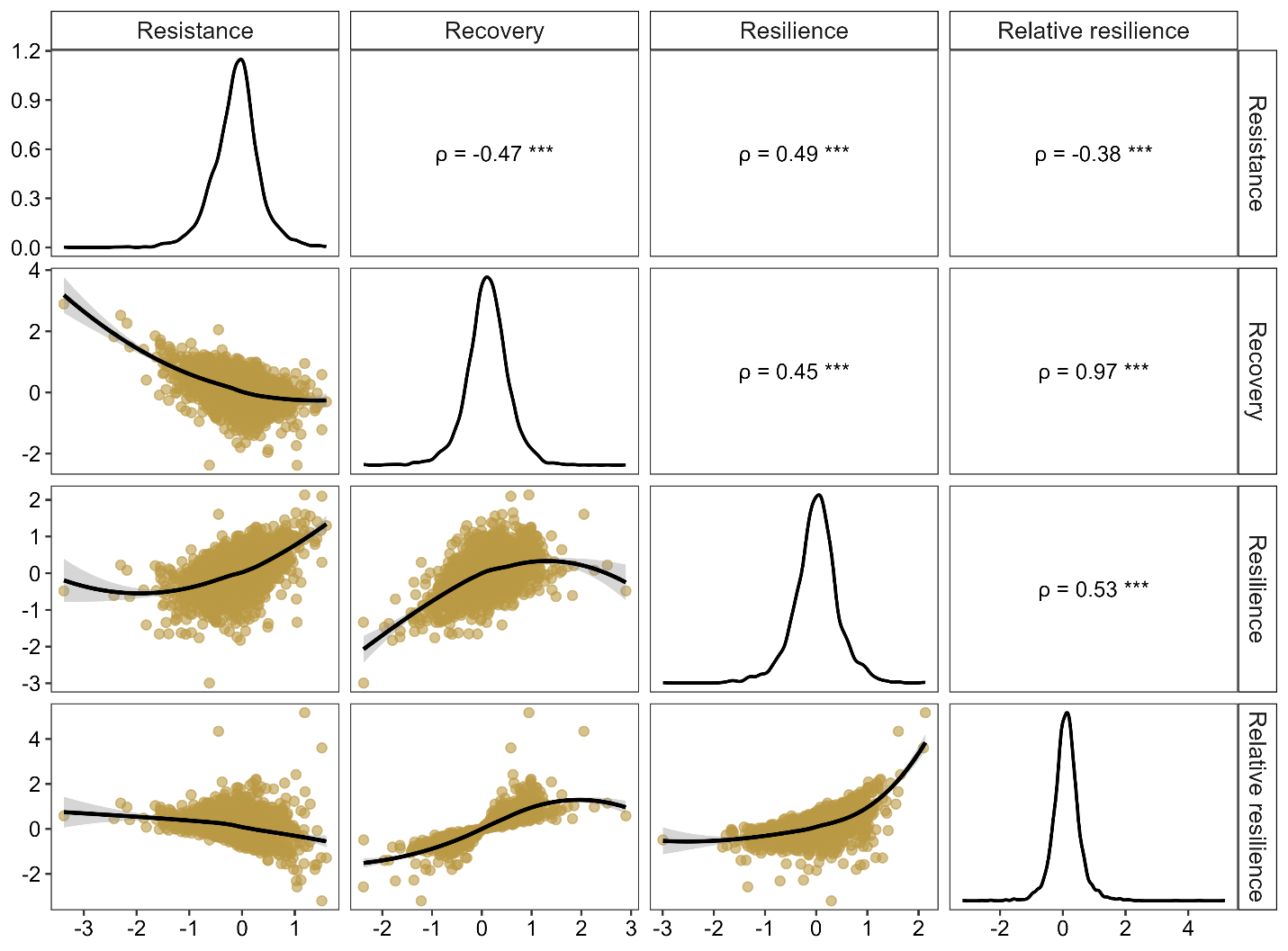

**Table S1**. Overview of FunDivEUROPE forest plot network.

|  | Finland | Germany | Italy | Poland | Romania | Spain |
| --- | --- | --- | --- | --- | --- | --- |
| Forest type | Boreal | Beech | Termophilous deciduous | Hemiboreal | Mountainous beech | Mediterranean mixed |
| Coordinates | 63 °N 30 °E | 51 °N 11 °E | 43 °N 11 °E | 53 °N 24 °E | 47 °N 26 °E | 41 °N 2 °W |
| Number of plots | 28 | 38 | 36 | 43 | 28 | 36 |
| Elevation range (m.a.s.l.) | 80–200 | 500–600 | 260–525 | 135–185 | 600–1000 | 960–1400 |
| Mean annual temperature (°C) | 2.1 | 6.8 | 13.0 | 6.9 | 6.8 | 10.2 |
| Mean annual precipitation (mm) | 700 | 775 | 850 | 625 | 800 | 500 |
| Drought events targeted | 2006 \| 1994 | 2003 \| 1976 | 2003 \| 1990 | 2002 \| 1992 | 2003 \| 2000 | 2005 \| 1990 |
| Summer CWD of the drought  year (mm) | -60.83 \| -39.00 | -62.14 \| -65.44 | -121.54 \| -107.14 | -70.60 \| -73.57 | -64.44 \| -53.79 | -157.75 \| -143.28 |
| Growing season CWD of the  drought year (mm) | -32.47 \| -22.57 | -39.27 \| -37.79 | -79.31 \| -65.44 | -51.26 \| -39.75 | -45.96 \| -33.21 | -116.98 \| -97.52 |
| Mean basal area (m^2^ ha ^-1^) | 22.8 | 35.7 | 27.2 | 37.8 | 51.3 | 22 |
| Species pool | *Betula pendula*  *Picea abies*  *Pinus sylvestris* | *Acer pseudoplatanus*  *Fagus sylvatica*  *Fraxinus excelsior*  *Picea abies*  *Quercus petraea* | *Castanea sativa*  *Ostrya carpinifolia*  *Quercus cerris*  *Quercus petraea*  *Quercus ilex* | *Betula pendula*  *Carpinus betulus*  *Picea abies*  *Pinus sylvestris*  *Quercus robur* | *Acer pseudoplatanus*  *Abies alba*  *Fagus sylvatica*  *Picea abies* | *Pinus nigra*  *Pinus sylvestris*  *Quercus faginea*  *Quercus ilex* |

**Table S2.** Descriptive statistics for the detrended tree ring chronologies of each study site from the FunDivEUROPE forest plot network.

| Country | N plots | N trees | N cores | Ranged used | EPS | Rbar | SSS | SNR | MS |
| --- | --- | --- | --- | --- | --- | --- | --- | --- | --- |
| Finland | 28 | 2086 | 442 | 1960-2011 | 0.983 | 0.150 | 0.924 | 56.257 | 0.21 |
| Italy | 36 | 2979 | 645 | 1902-2012 | 0.989 | 0.136 | 0.795 | 86.333 | 0.24 |
| Germany | 38 | 1793 | 524 | 1779-2011 | 0.978 | 0.084 | 0.731 | 45.302 | 0.29 |
| Poland | 43 | 2083 | 441 | 1713-2011 | 0.976 | 0.089 | 0.732 | 41.110 | 0.27 |
| Romania | 28 | 1350 | 399 | 1740-2011 | 0.978 | 0.101 | 0.578 | 44.332 | 0.26 |
| Spain | 36 | 2648 | 458 | 1821-2011 | 0.989 | 0.183 | 0.874 | 90.971 | 0.28 |

Notes. EPS: Expressed population signal; Rbar: mean inter-series correlation; SSS: subsample signal strength; SNR: signal-to-noise ratio; MS: mean sensitivity.

**Table S3.** Pearson’s correlation (*r*) between tree diameter (DBH) with height (H) and age.

| Species code | Full species name | DBH-H | DBH-Age |
| --- | --- | --- | --- |
| ABIALB | *Abies alba* | 0.865 | 0.305 |
| ACEPSE | *Acer pseudoplatanus* | 0.707 | 0.627 |
| BETPEN | *Betula pendula* | 0.892 | 0.566 |
| CARBET | *Carpinus betulus* | 0.766 | 0.437 |
| CASSAT | *Castanea sativa* | 0.596 | 0.643 |
| FAGSYL | *Fagus sylvatica* | 0.837 | 0.577 |
| FRAEXC | *Fraxinus excelsior* | 0.703 | 0.639 |
| OSTCAR | *Ostrya carpinifolia* | 0.682 | 0.388 |
| PICABI | *Picea abies* | 0.892 | 0.606 |
| PINSYL | *Pinus sylvestris* | 0.600 | 0.633 |
| PINNIG | *Pinus nigra* | 0.676 | 0.607 |
| QUECER | *Quercus cerris* | 0.726 | 0.407 |
| QUEILE | *Quercus ilex* | 0.613 | 0.249 |
| QUEFAG | *Quercus faginea* | 0.705 | 0.349 |
| QUEPET | *Quercus petraea* | 0.825 | 0.798 |
| QUEROB | *Quercus robur* | 0.719 | 0.613 |

**Table S4. Model results for tree resistance (Rt) in response to the recent drought.** Draws were sampled using sampling (NUTS). For each model parameter: β represents the standardised regression estimate, SE the standard error, Bulk_ESS and Tail_ESS the effective sample size measures. Rhat is the potential scale reduction factor on split chains (at convergence, Rhat = 1), and the 95% CI are the lower (left) and upper (right) credible intervals, respectively. The number of observations is 2909. Nonzero CIs are highlighted in bold blue.

| Multilevel Hyperparameters: | | | | | | | | |
| --- | --- | --- | --- | --- | --- | --- | --- | --- |
| Random effects: |  |  |  |  |  |  | | |
| Plot (208 levels) | **β** | **SE** | **Rhat** | **Bulk_ESS** | **Tail_ESS** | **95% CI** | | |
| Sd(Intercept) | **0.14** | **0.01** | **1.00** | **1581** | **2360** | **0.12** | **0.17** | |
| Sd(σ Intercept) | **0.26** | **0.02** | **1.00** | **1566** | **2321** | **0.22** | **0.31** | |
| Species (16 levels) | **β** | **SE** | **Rhat** | **Bulk_ESS** | **Tail_ESS** | **95% CI** | | |
| Sd(Intercept) | **0.21** | **0.08** | **1.00** | **1104** | **811** | **0.08** | **0.37** | |
| Sd(PreDr10) | 0.03 | 0.02 | 1.00 | 1442 | 2240 | 0.00 | 0.09 | |
| Sd(Height) | 0.03 | 0.02 | 1.00 | 2401 | 2568 | 0.00 | 0.09 | |
| Sd(LCR) | 0.02 | 0.02 | 1.00 | 1981 | 2623 | 0.00 | 0.06 | |
| Sd(Age) | 0.04 | 0.02 | 1.00 | 1339 | 1580 | 0.00 | 0.09 | |
| Sd(SR) | 0.02 | 0.01 | 1.00 | 1893 | 1848 | 0.00 | 0.06 | |
| Sd(BAT subplot) | 0.02 | 0.02 | 1.00 | 2053 | 1997 | 0.00 | 0.06 | |
| Sd(CWD_gs) | 0.08 | 0.07 | 1.00 | 1271 | 1699 | 0.00 | 0.26 | |
| Sd(σ Intercept) | **0.45** | **0.09** | **1.01** | **1474** | **2417** | **0.31** | **0.67** | |
| Species:Country (26 levels) | **β** | **SE** | **Rhat** | **Bulk_ESS** | **Tail_ESS** | **95% CI** | | |
| Sd(Intercept) | **0.09** | **0.06** | **1.00** | **855** | **1169** | **0.01** | | **0.23** |
| Sd(PreDr10) | 0.03 | 0.02 | 1.00 | 1098 | 1625 | 0.00 | | 0.08 |
| Sd(Height) | 0.03 | 0.02 | 1.01 | 1490 | 1865 | 0.00 | | 0.09 |
| Sd(LCR) | 0.02 | 0.02 | 1.00 | 1703 | 1966 | 0.00 | | 0.06 |
| Sd(Age) | 0.04 | 0.03 | 1.00 | 1258 | 1610 | 0.00 | | 0.10 |
| Sd(SR) | 0.03 | 0.02 | 1.00 | 1334 | 1595 | 0.00 | | 0.07 |
| Sd(BAT subplot) | 0.03 | 0.02 | 1.00 | 1454 | 1936 | 0.00 | | 0.07 |
| Sd(CWD_gs) | **0.10** | **0.07** | **1.00** | **931** | **2182** | **0.01** | | **0.25** |
| Fixed effects: |  |  |  |  |  |  | |  |
| Variable | **β** | **SE** | **Rhat** | **Bulk_ESS** | **Tail_ESS** | **95% CI** | | |
| Intercept | 0.02 | 0.07 | 1.00 | 1875 | 2482 | -0.11 | | 0.16 |
| σ intercept | **-0.56** | **0.11** | **1.00** | **633** | **1234** | **-0.79** | | **-0.34** |
| PreDr10 | -0.02 | 0.02 | 1.00 | 2613 | 2414 | -0.07 | | 0.01 |
| Height | 0.03 | 0.03 | 1.00 | 3366 | 2812 | -0.02 | | 0.08 |
| LCR | 0.01 | 0.01 | 1.00 | 3234 | 2879 | -0.02 | | 0.04 |
| Age | 0.02 | 0.02 | 1.00 | 3257 | 3109 | -0.02 | | 0.07 |
| SR | -0.00 | 0.02 | 1.00 | 2470 | 2580 | -0.04 | | 0.03 |
| BAT subplot | -0.03 | 0.02 | 1.00 | 3075 | 2836 | -0.06 | | 0.00 |
| CWD_gs | **0.19** | **0.07** | **1.00** | **2034** | **1826** | **0.04** | | **0.31** |

**Table S5. Model results for tree recovery (Rc) in response to the recent drought.** Draws were sampled using sampling (NUTS). For each model parameter: β represents the standardised regression estimate, SE the standard error, Bulk_ESS and Tail_ESS are effective sample size measures. Rhat is the potential scale reduction factor on split chains (at convergence, Rhat = 1), and the 95% CI are the lower (left) and upper (right) credible intervals, respectively. The number of observations is 2909. Nonzero CIs are highlighted in bold blue.

| Multilevel Hyperparameters: | | | | | | | | |
| --- | --- | --- | --- | --- | --- | --- | --- | --- |
| Random effects: |  |  |  |  |  |  | | |
| Plot (208 levels) | **β** | **SE** | **Rhat** | **Bulk_ESS** | **Tail_ESS** | **95% CI** | | |
| Sd(Intercept) | **0.10** | **0.01** | **1.00** | **1180** | **1653** | **0.08** | **0.13** | |
| Sd(σ Intercept) | **0.35** | **0.02** | **1.01** | **1360** | **2114** | **0.31** | **0.40** | |
| Species (16 levels) | **β** | **SE** | **Rhat** | **Bulk_ESS** | **Tail_ESS** | **95% CI** | | |
| Sd(Intercept) | **0.14** | **0.10** | **1.00** | **719** | **1498** | **0.01** | **0.37** | |
| Sd(PreDr10) | 0.03 | 0.02 | 1.00 | 1506 | 2095 | 0.00 | 0.07 | |
| Sd(Height) | 0.04 | 0.03 | 1.01 | 1566 | 1889 | 0.00 | 0.11 | |
| Sd(LCR) | 0.05 | 0.03 | 1.00 | 1048 | 1189 | 0.00 | 0.10 | |
| Sd(Age) | 0.04 | 0.03 | 1.00 | 1253 | 1796 | 0.00 | 0.10 | |
| Sd(SR) | 0.02 | 0.02 | 1.00 | 1718 | 1771 | 0.00 | 0.06 | |
| Sd(BAT subplot) | 0.02 | 0.02 | 1.00 | 1612 | 1854 | 0.00 | 0.06 | |
| Sd(CWD_gs) | 0.13 | 0.10 | 1.00 | 1212 | 1748 | 0.00 | 0.37 | |
| Sd(σ Intercept) | **0.37** | **0.08** | **1.00** | **1175** | **1623** | **0.25** | **0.56** | |
| Species:Country (26 levels) | **β** | **SE** | **Rhat** | **Bulk_ESS** | **Tail_ESS** | **95% CI** | | |
| Sd(Intercept) | **0.29** | **0.07** | **1.00** | **1506** | **1752** | **0.17** | | **0.43** |
| Sd(PreDr10) | 0.03 | 0.02 | 1.00 | 1244 | 1808 | 0.00 | | 0.07 |
| Sd(Height) | 0.06 | 0.03 | 1.00 | 1129 | 1151 | 0.00 | | 0.13 |
| Sd(LCR) | 0.03 | 0.02 | 1.00 | 1157 | 2230 | 0.00 | | 0.08 |
| Sd(Age) | **0.06** | **0.03** | **1.00** | **921** | **1215** | **0.01** | | **0.13** |
| Sd(SR) | 0.03 | 0.02 | 1.01 | 921 | 1105 | 0.00 | | 0.08 |
| Sd(BAT subplot) | 0.02 | 0.01 | 1.00 | 1460 | 1894 | 0.00 | | 0.05 |
| Sd(CWD_gs) | 0.11 | 0.08 | 1.00 | 1199 | 1705 | 0.00 | | 031 |
| Fixed effects: |  |  |  |  |  |  | |  |
| Variable | **β** | **SE** | **Rhat** | **Bulk_ESS** | **Tail_ESS** | **95% CI** | | |
| Intercept | -0.06 | 0.08 | 1.00 | 1995 | 2530 | -0.22 | | 0.11 |
| σ intercept | **-0.65** | **0.10** | **1.00** | **547** | **990** | **-0.84** | | **-0.45** |
| PreDr10 | 0.03 | 0.02 | 1.00 | 2876 | 2618 | -0.01 | | 0.06 |
| Height | -0.03 | 0.03 | 1.00 | 2713 | 2964 | -0.09 | | 0.02 |
| LCR | 0.03 | 0.02 | 1.00 | 2887 | 3015 | -0.01 | | 0.06 |
| Age | -0.03 | 0.03 | 1.00 | 2974 | 2651 | -0.09 | | 0.02 |
| SR | 0.01 | 0.02 | 1.00 | 2707 | 2774 | -0.02 | | 0.05 |
| BAT subplot | 0.01 | 0.01 | 1.00 | 3319 | 3312 | -0.02 | | 0.04 |
| CWD_gs | -0.12 | 0.09 | 1.00 | 1831 | 2457 | -0.29 | | 0.06 |

**Table S6. Model results for tree resilience (Rs) in response to the recent drought.** Draws were sampled using sampling (NUTS). For each model parameter: β represents the standardised regression estimate, SE the standard error, Bulk_ESS and Tail_ESS are effective sample size measures. Rhat is the potential scale reduction factor on split chains (at convergence, Rhat = 1), and the 95% CI are the lower (left) and upper (right) credible intervals, respectively. The number of observations is 2909. Nonzero CIs are highlighted in bold blue.

| Multilevel Hyperparameters: | | | | | | | | |
| --- | --- | --- | --- | --- | --- | --- | --- | --- |
| Random effects: |  |  |  |  |  |  | | |
| Plot (208 levels) | **β** | **SE** | **Rhat** | **Bulk_ESS** | **Tail_ESS** | **95% CI** | | |
| Sd(Intercept) | **0.14** | **0.02** | **1.00** | **1227** | **2037** | **0.11** | **0.17** | |
| Sd(σ Intercept) | **0.26** | **0.02** | **1.00** | **1614** | **2555** | **0.23** | **0.31** | |
| Species (16 levels) | **β** | **SE** | **Rhat** | **Bulk_ESS** | **Tail_ESS** | **95% CI** | | |
| Sd(Intercept) | **0.16** | **0.12** | **1.00** | **925** | **1725** | **0.01** | **0.44** | |
| Sd(PreDr10) | 0.06 | 0.05 | 1.00 | 731 | 1867 | 0.00 | 0.18 | |
| Sd(Height) | 0.05 | 0.04 | 1.00 | 1361 | 1413 | 0.00 | 0.14 | |
| Sd(LCR) | 0.02 | 0.02 | 1.00 | 1690 | 1721 | 0.00 | 0.06 | |
| Sd(Age) | 0.03 | 0.02 | 1.00 | 2025 | 2084 | 0.00 | 0.08 | |
| Sd(SR) | 0.04 | 0.02 | 1.00 | 1187 | 1756 | 0.00 | 0.08 | |
| Sd(BAT subplot) | 0.02 | 0.02 | 1.00 | 1987 | 2354 | 0.00 | 0.06 | |
| CWD_gs | **0.20** | **0.15** | **1.00** | **847** | **1316** | **0.01** | **0.57** | |
| Sd(σ Intercept) | **0.40** | **0.08** | **1.00** | **866** | **1769** | **0.27** | **0.59** | |
| Species: Country (26 levels) | **β** | **SE** | **Rhat** | **Bulk_ESS** | **Tail_ESS** | **95% CI** | | |
| Sd(Intercept) | **0.33** | **0.09** | **1.00** | **1813** | **2324** | **0.19** | | **0.54** |
| Sd(PreDr10) | **0.08** | **0.03** | **1.01** | **567** | **462** | **0.02** | | **0.15** |
| Sd(Height) | **0.10** | **0.04** | **1.00** | **954** | **816** | **0.02** | | **0.19** |
| Sd(LCR) | 0.02 | 0.02 | 1.00 | 2080 | 1769 | 0.00 | | 0.06 |
| Sd(Age) | 0.04 | 0.02 | 1.00 | 1246 | 1705 | 0.00 | | 0.09 |
| Sd(SR) | 0.03 | 0.02 | 1.00 | 1332 | 1918 | 0.00 | | 0.07 |
| Sd(BAT subplot) | 0.02 | 0.02 | 1.00 | 1395 | 1506 | 0.00 | | 0.06 |
| CWD_gs | **0.20** | **0.14** | **1.00** | **771** | **1476** | **0.01** | | **0.52** |
| Fixed effects: |  |  |  |  |  |  | |  |
| Variable | **β** | **SE** | **Rhat** | **Bulk_ESS** | **Tail_ESS** | **95% CI** | | |
| Intercept | -0.01 | 0.10 | 1.00 | 1832 | 2146 | -0.21 | | 0.21 |
| σ intercept | **-0.49** | **0.10** | **1.01** | **504** | **1022** | **-0.69** | | **-0.28** |
| PreDr10 | -0.01 | 0.03 | 1.00 | 1980 | 2435 | -0.08 | | 0.06 |
| Height | -0.00 | 0.04 | 1.00 | 2723 | 2445 | -0.08 | | 0.07 |
| LCR | **0.04** | **0.02** | **1.00** | **3615** | **3085** | **0.01** | | **0.07** |
| Age | -0.02 | 0.02 | 1.00 | 3501 | 3264 | -0.07 | | 0.02 |
| SR | 0.01 | 0.02 | 1.00 | 2662 | 3017 | -0.03 | | 0.05 |
| BAT subplot | 0.00 | 0.02 | 1.00 | 3517 | 2650 | -0.03 | | 0.04 |
| CWD_gs | 0.04 | 0.12 | 1.00 | 2008 | 2351 | -0.24 | | 0.27 |

**Table S7. Model results for tree relative resilience (RRs) in response to the recent drought.** Draws were sampled using sampling (NUTS). For each model parameter: β represents the standardised regression estimate, SE the standard error, Bulk_ESS and Tail_ESS are effective sample size measures. Rhat is the potential scale reduction factor on split chains (at convergence, Rhat = 1), and the 95% CI are the lower (left) and upper (right) credible intervals, respectively. The number of observations is 2909. Nonzero CIs are highlighted in bold blue.

| Multilevel Hyperparameters: | | | | | | | | |
| --- | --- | --- | --- | --- | --- | --- | --- | --- |
| Random effects: |  |  |  |  |  |  | | |
| Plot (208 levels) | **β** | **SE** | **Rhat** | **Bulk_ESS** | **Tail_ESS** | **95% CI** | | |
| Sd(Intercept) | **0.15** | **0.02** | **1.00** | **1501** | **2075** | **0.11** | **0.19** | |
| Sd(σ Intercept) | **0.33** | **0.02** | **1.00** | **1731** | **2899** | **0.29** | **0.38** | |
| Species (16 levels) | **β** | **SE** | **Rhat** | **Bulk_ESS** | **Tail_ESS** | **95% CI** | | |
| Sd(Intercept) | **0.18** | **0.13** | **1.00** | **1224** | **2329** | **0.01** | **0.47** | |
| Sd(PreDr10) | 0.04 | 0.03 | 1.00 | 1730 | 2361 | 0.00 | 0.11 | |
| Sd(Height) | 0.05 | 0.04 | 1.00 | 2554 | 2348 | 0.00 | 0.16 | |
| Sd(LCR) | **0.06** | **0.03** | **1.00** | **1369** | **1729** | **0.01** | **0.14** | |
| Sd(Age) | 0.05 | 0.03 | 1.00 | 1878 | 2645 | 0.00 | 0.13 | |
| Sd(SR) | 0.04 | 0.03 | 1.00 | 2021 | 2086 | 0.00 | 0.10 | |
| Sd(BAT subplot) | 0.03 | 0.02 | 1.00 | 2340 | 2495 | 0.00 | 0.08 | |
| CWD_gs | **0.14** | **0.12** | **1.00** | **1727** | **2308** | **0.01** | **0.42** | |
| Sd(σ Intercept) | **0.41** | **0.08** | **1.00** | **1240** | **2301** | **0.28** | **0.60** | |
| Species:Country (26 levels) | **β** | **SE** | **Rhat** | **Bulk_ESS** | **Tail_ESS** | **95% CI** | | |
| Sd(Intercept) | **0.39** | **0.09** | **1.00** | **2532** | **2428** | **0.23** | | **0.59** |
| Sd(PreDr10) | 0.05 | 0.03 | 1.00 | 1554 | 1662 | 0.00 | | 0.11 |
| Sd(Height) | **0.11** | **0.05** | **1.00** | **1149** | **1086** | **0.02** | | **0.21** |
| Sd(LCR) | 0.05 | 0.03 | 1.00 | 1216 | 2466 | 0.00 | | 0.11 |
| Sd(Age) | 0.07 | 0.04 | 1.00 | 1241 | 1763 | 0.00 | | 0.16 |
| Sd(SR) | 0.05 | 0.03 | 1.00 | 1109 | 1152 | 0.00 | | 0.11 |
| Sd(BAT subplot) | 0.03 | 0.02 | 1.00 | 1648 | 2159 | 0.00 | | 0.08 |
| CWD_gs | 0.13 | 0.10 | 1.00 | 1626 | 2380 | 0.00 | | 0.37 |
| Fixed effects: |  |  |  |  |  |  | |  |
| Variable | **β** | **SE** | **Rhat** | **Bulk_ESS** | **Tail_ESS** | **95% CI** | | |
| Intercept | -0.01 | 0.11 | 1.00 | 2325 | 2274 | -0.21 | | 0.20 |
| σ intercept | **-0.41** | **0.10** | **1.00** | **1158** | **1933** | **-0.61** | | **-0.20** |
| PreDr10 | 0.03 | 0.03 | 1.00 | 3935 | 2830 | -0.03 | | 0.08 |
| Height | -0.05 | 0.05 | 1.00 | 3448 | 3142 | -0.14 | | 0.04 |
| LCR | 0.04 | 0.03 | 1.00 | 4054 | 3303 | -0.01 | | 0.10 |
| Age | -0.05 | 0.03 | 1.00 | 4231 | 3113 | -0.12 | | 0.02 |
| SR | 0.02 | 0.03 | 1.00 | 3813 | 2711 | -0.03 | | 0.07 |
| BAT subplot | 0.02 | 0.02 | 1.00 | 4887 | 2895 | -0.03 | | 0.06 |
| CWD_gs | -0.13 | 0.11 | 1.00 | 3251 | 3140 | -0.34 | | 0.09 |

**Table S8.** Model results for each resilience model. For each model, the marginal and conditional *R*^2^ as well as the 95% credible intervals (CIs) are shown.

| Model | Marginal *R*^2^ | 95% CI | Conditional *R*^2^ | 95% CI |
| --- | --- | --- | --- | --- |
| Resistance (Rt) | 0.043 | 0.003, 0.088 | 0.128 | 0.111, 0.145 |
| Recovery (Rc) | 0.026 | 0.001, 0.081 | 0.147 | 0.131, 0.164 |
| Resilience (Rs) | 0.013 | 0.000, 0.061 | 0.145 | 0.125, 0.167 |
| Rel. resilience (RRs) | 0.036 | 0.003, 0.108 | 0.208 | 0.186, 0.231 |

**Table caption illustrating the intraspecific effect of tree height on Lloret's resilience components:**

Summary of linear mixed models (LMM) computed using R-package (*lm4*) testing within-species (*N*=16) effects of predrought growth (PreDr_growth), tree height, tree age, basal area of trees taller than the cored trees in the subplot (BAT subplot), live crown ratio, species richness and slope as fixed effects and forest plot as a random intercept term on four components of individual tree-level resilience to drought: resistance (Rt), recovery (Rc), resilience (Rs) and relative resilience (RRs). The tables presented here consist of individual LMMs computed for each species. Significant effects (*p* <0.05) are shown in bold. Statistics show the standardised regression estimate (β), standard error (SE), t-value, degrees of freedom (df), *p*-value, and 95% of credible interval (CI). SD is the estimated standard deviation of the forest plots within each country. R² LMM(M) and R² LMM(C) represent the explained variation of fixed and fixed plus random factors, respectively.

**Table S9.** Dependent variable: resilience (Rs), tree species: *Abies alba* from Romania

| Random effects | Name | Variance | *SD* |  |  |  |  |
| --- | --- | --- | --- | --- | --- | --- | --- |
| Plot | Intercept | 0.048 | 0.220 |  |  |  |  |
| Residual |  | 0.891 | 0.944 |  |  |  |  |
| Fixed effects |  |  |  |  |  |  |  |
| Variable | **β** | ***SE*** | ***t-value*** | ***df*** | ***P*-value** | **95% CI** | |
| Slope | 0.190 | 0.252 | 0.753 | 5.992 | 0.480 | -0.426 | 0.805 |
| Species richness | 0.213 | 0.147 | 1.455 | 6.886 | 0.190 | -0.134 | 0.561 |
| Live crown ratio | -0.044 | 0.094 | -0.467 | 68.454 | 0.642 | -0.230 | 0.143 |
| Age | -0.006 | 0.116 | -0.054 | 90.155 | 0.957 | -0.237 | 0.224 |
| Height | **-0.201** | **0.096** | **-2.088** | **90.706** | **0.040** | **-0.393** | **-0.010** |
| PreDr_ growth | -0.175 | 0.119 | -1.469 | 85.624 | 0.145 | -0.411 | 0.062 |
| Intercept | **0.532** | **0.207** | **2.568** | **7.443** | **0.035** | **0.048** | **1.016** |
| R^2^ _LMM (C)_ | 0.174 |  |  |  |  |  |  |
| R^2^ _LMM (M)_ | 0.129 |  |  |  |  |  |  |

**Table S10.** Dependent variable: relative resilience (RRs), tree species: *Abies alba* from Romania

| Random effects | Name | Variance | *SD* |  |  |  |  |
| --- | --- | --- | --- | --- | --- | --- | --- |
| Plot | Intercept | 0.044 | 0.210 |  |  |  |  |
| Residual |  | 0.566 | 0.752 |  |  |  |  |
| Fixed effects |  |  |  |  |  |  |  |
| Variable | **β** | ***SE*** | ***t-value*** | ***df*** | ***P*-value** | **95% CI** | |
| Slope | 0.141 | 0.212 | 0.666 | 7.028 | 0.527 | -0.360 | 0.643 |
| Species richness | 0.043 | 0.123 | 0.351 | 8.034 | 0.735 | -0.241 | 0.327 |
| Live crown ratio | -0.047 | 0.076 | -0.624 | 76.600 | 0.534 | -0.198 | 0.103 |
| Age | 0.055 | 0.093 | 0.595 | 90.931 | 0.553 | -0.130 | 0.241 |
| Height | **-0.228** | **0.077** | **-2.947** | **90.315** | **0.004** | **-0.381** | **-0.074** |
| PreDr_growth | -0.043 | 0.095 | -0.448 | 85.829 | 0.655 | -0.231 | 0.146 |
| Intercept | **0.874** | **0.174** | **5.024** | **8.540** | **0.001** | **0.477** | **1.270** |
| R^2^ _LMM (C)_ | 0.179 |  |  |  |  |  |  |
| R^2^ _LMM (M)_ | 0.115 |  |  |  |  |  |  |

**Table S11.** Dependent variable: recovery (Rc), tree species: *Abies alba* from Romania

| Random effects | Name | Variance | *SD* |  |  |  |  |
| --- | --- | --- | --- | --- | --- | --- | --- |
| Plot | Intercept | 0.088 | 0.296 |  |  |  |  |
| Residual |  | 0.467 | 0.684 |  |  |  |  |
| Fixed effects |  |  |  |  |  |  |  |
| Variable | **β** | ***SE*** | ***t-value*** | ***df*** | ***P*-value** | **95% CI** | |
| Slope | 0.131 | 0.237 | 0.553 | 9.894 | 0.593 | -0.398 | 0.660 |
| Species richness | -0.143 | 0.136 | -1.051 | 11.009 | 0.316 | -0.442 | 0.156 |
| Live crown ratio | -0.067 | 0.071 | -0.933 | 89.261 | 0.353 | -0.208 | 0.075 |
| Age | 0.024 | 0.086 | 0.275 | 89.823 | 0.784 | -0.148 | 0.195 |
| Height | **-0.221** | **0.071** | **-3.098** | **88.723** | **0.003** | **-0.362** | **-0.079** |
| PreDr_growth | 0.016 | 0.087 | 0.179 | 85.737 | 0.858 | -0.157 | 0.189 |
| Intercept | **0.672** | **0.191** | **3.515** | **11.280** | **0.005** | **0.252** | **1.091** |
| R^2^ _LMM (C)_ | 0.259 |  |  |  |  |  |  |
| R^2^ _LMM (M)_ | 0.119 |  |  |  |  |  |  |

**Table S12.** Dependent variable: resilience (Rs), tree species: *Betula pendula* from Finland

| Random effects | Name | Variance | *SD* |  |  |  |  |
| --- | --- | --- | --- | --- | --- | --- | --- |
| Plot | Intercept | 0.054 | 0.234 |  |  |  |  |
| Residual |  | 0.305 | 0.553 |  |  |  |  |
| Fixed effects |  |  |  |  |  |  |  |
| Variable | **β** | ***SE*** | ***t-value*** | ***df*** | ***P-value*** | **95% CI** | |
| Slope | 0.082 | 0.104 | 0.790 | 11.260 | 0.446 | -0.146 | 0.309 |
| Species richness | 0.073 | 0.134 | 0.546 | 11.267 | 0.596 | -0.220 | 0.366 |
| Live crown ratio | 0.123 | 0.091 | 1.349 | 129.634 | 0.180 | -0.058 | 0.304 |
| BAT subplot | 0.216 | 0.141 | 1.529 | 114.826 | 0.129 | -0.064 | 0.497 |
| Age | 0.365 | 0.424 | 0.861 | 68.739 | 0.392 | -0.480 | 1.210 |
| Height | **0.547** | **0.225** | **2.432** | **78.871** | **0.017** | **0.099** | **0.994** |
| PreDr_growth | **-0.708** | **0.286** | **-2.474** | **126.841** | **0.015** | **-1.273** | **-0.142** |
| Intercept | 0.522 | 0.379 | 1.378 | 42.619 | 0.175 | -0.242 | 1.286 |
| R^2^ _LMM (C)_ | 0.307 |  |  |  |  |  |  |
| R^2^ _LMM (M)_ | 0.184 |  |  |  |  |  |  |

**Table S13.** Dependent variable: relative resilience (RRs), tree species: *Carpinus betulus* from Poland

| Random effects | Name | Variance | *SD* |  |  |  |  |
| --- | --- | --- | --- | --- | --- | --- | --- |
| Plot | Intercept | 0.000 | 0.000 |  |  |  |  |
| Residual |  | 1.137 | 1.066 |  |  |  |  |
| Fixed effects |  |  |  |  |  |  |  |
| Variable | **β** | ***SE*** | ***t-value*** | ***df*** | ***P-value*** | **95% CI** | |
| Species richness | 0.164 | 0.141 | 1.165 | 63 | 0.248 | -0.118 | 0.446 |
| Live crown ratio | **-0.362** | **0.163** | **-2.224** | **63** | **0.030** | **-0.688** | **-0.037** |
| BAT subplot | -0.068 | 0.144 | -0.471 | 63 | 0.640 | -0.355 | 0.220 |
| Age | -0.094 | 0.222 | -0.425 | 63 | 0.672 | -0.538 | 0.350 |
| Height | **-0.915** | **0.394** | **-2.322** | **63** | **0.023** | **-1.703** | **-0.128** |
| PreDr_growth | 0.068 | 0.217 | 0.311 | 63 | 0.756 | -0.366 | 0.501 |
| Intercept | -0.132 | 0.227 | -0.582 | 63 | 0.562 | -0.587 | 0.322 |
| R^2^ _LMM (C)_ | NA |  |  |  |  |  |  |
| R^2^ _LMM (M)_ | 0.155 |  |  |  |  |  |  |

**Table S14.** Dependent variable: recovery (Rc), tree species: *Carpinus betulus* from Poland

| Random effects | Name | Variance | *SD* |  |  |  |  |
| --- | --- | --- | --- | --- | --- | --- | --- |
| Plot | Intercept | 0.000 | 0.000 |  |  |  |  |
| Residual |  | 0.855 | 0.925 |  |  |  |  |
| Fixed effects |  |  |  |  |  |  |  |
| Variable | **β** | ***SE*** | ***t-value*** | ***df*** | ***P-value*** | **95% CI** | |
| Species richness | 0.208 | 0.122 | 1.701 | 63 | 0.0938 | -0.0363 | 0.4529 |
| Live crown ratio | -0.280 | 0.141 | -1.980 | 63 | 0.0521 | -0.5621 | 0.0026 |
| BAT subplot | -0.076 | 0.125 | -0.608 | 63 | 0.5457 | -0.3251 | 0.1735 |
| Age | 0.089 | 0.193 | 0.464 | 63 | 0.6443 | -0.2957 | 0.4744 |
| Height | **-0.690** | **0.342** | **-2.019** | **63** | **0.0477** | **-1.3732** | **-0.0071** |
| PreDr_growth | -0.011 | 0.188 | -0.059 | 63 | 0.9532 | -0.3870 | 0.3648 |
| Intercept | -0.069 | 0.197 | -0.350 | 63 | 0.7273 | -0.4630 | 0.3249 |
| R^2^ _LMM (C)_ | NA |  |  |  |  |  |  |
| R^2^ _LMM (M)_ | 0.114 |  |  |  |  |  |  |

**Table S15.** Dependent variable: resilience (Rs), tree species: *Fagus sylvatica* from Germany

| Random effects | Name | Variance | *SD* |  |  |  |  |
| --- | --- | --- | --- | --- | --- | --- | --- |
| Plot | Intercept | 0.161 | 0.401 |  |  |  |  |
| Residual |  | 1.272 | 1.128 |  |  |  |  |
| Fixed effects |  |  |  |  |  |  |  |
| Variable | **β** | ***SE*** | ***t-value*** | ***df*** | ***P-value*** | **95% CI** | |
| Slope | -0.107 | 0.111 | -0.959 | 35.065 | 0.344 | -0.332 | 0.119 |
| Species richness | 0.053 | 0.138 | 0.384 | 24.975 | 0.704 | -0.232 | 0.338 |
| Live crown ratio | 0.05 | 0.104 | 0.475 | 191.389 | 0.635 | -0.156 | 0.256 |
| BAT subplot | 0.008 | 0.072 | 0.114 | 188.428 | 0.909 | -0.134 | 0.15 |
| Age | -0.117 | 0.115 | -1.018 | 173.084 | 0.31 | -0.345 | 0.11 |
| Height | **-0.325** | **0.132** | **-2.469** | **183.67** | **0.014** | **-0.585** | **-0.065** |
| PreDr_growth | -0.022 | 0.025 | -0.881 | 186.892 | 0.38 | -0.07 | 0.027 |
| Intercept | -0.051 | 0.147 | -0.345 | 53.23 | 0.731 | -0.345 | 0.244 |
| R^2^ _LMM (C)_ | 0.200 |  |  |  |  |  |  |
| R^2^ _LMM (M)_ | 0.099 |  |  |  |  |  |  |

**Table S16.** Dependent variable: recovery (Rc), tree species: *Fagus sylvatica* from Germany

| Random effects | Name | Variance | *SD* |  |  |  |  |
| --- | --- | --- | --- | --- | --- | --- | --- |
| Plot | Intercept | 0.000 | 0.000 |  |  |  |  |
| Residual |  | 0.611 | 0.782 |  |  |  |  |
| Fixed effects |  |  |  |  |  |  |  |
| Variable | **β** | ***SE*** | ***t-value*** | ***df*** | ***P-value*** | **95% CI** | |
| Slope | -0.061 | 0.058 | -1.055 | 192 | 0.293 | -0.175 | 0.053 |
| Species richness | 0.131 | 0.067 | 1.96 | 192 | 0.051 | -0.001 | 0.262 |
| Live crown ratio | 0.101 | 0.068 | 1.487 | 192 | 0.139 | -0.033 | 0.235 |
| BAT subplot | 0.033 | 0.046 | 0.711 | 192 | 0.478 | -0.058 | 0.124 |
| Age | -0.021 | 0.072 | -0.29 | 192 | 0.772 | -0.164 | 0.122 |
| Height | **-0.224** | **0.084** | **-2.665** | **192** | **0.008** | **-0.39** | **-0.058** |
| PreDr_growth | 0.008 | 0.017 | 0.503 | 192 | 0.615 | -0.024 | 0.041 |
| Intercept | **-0.234** | **0.082** | **-2.863** | **192** | **0.005** | **-0.395** | **-0.073** |
| R^2^ _LMM (C)_ | NA |  |  |  |  |  |  |
| R^2^ _LMM (M)_ | 0.137 |  |  |  |  |  |  |

**Table S17.** Dependent variable: resilience (Rs), tree species: *Fraxinus excelsior* from Germany

| Random effects | Name | Variance | *SD* |  |  |  |  |
| --- | --- | --- | --- | --- | --- | --- | --- |
| Plot | Intercept | 0.000 | 0.000 |  |  |  |  |
| Residual |  | 0.221 | 0.471 |  |  |  |  |
| Fixed effects |  |  |  |  |  |  |  |
| Variable | **β** | ***SE*** | ***t-value*** | ***df*** | ***P-value*** | **95% CI** | |
| Slope | 0.027 | 0.048 | 0.566 | 122 | 0.572 | -0.067 | 0.121 |
| Species richness | -0.040 | 0.055 | -0.729 | 122 | 0.468 | -0.148 | 0.068 |
| Live crown ratio | 0.086 | 0.058 | 1.501 | 122 | 0.136 | -0.028 | 0.200 |
| BAT subplot | -0.033 | 0.043 | -0.767 | 122 | 0.444 | -0.117 | 0.052 |
| Age | 0.033 | 0.049 | 0.660 | 122 | 0.510 | -0.065 | 0.131 |
| Height | **-0.162** | **0.067** | **-2.427** | **122** | **0.017** | **-0.293** | **-0.030** |
| PreDr_ growth | **-0.173** | **0.074** | **-2.345** | **122** | **0.021** | **-0.320** | **-0.027** |
| Intercept | -0.019 | 0.087 | -0.214 | 122 | 0.831 | -0.190 | 0.153 |
| R^2^ _LMM (C)_ | NA |  |  |  |  |  |  |
| R^2^ _LMM (M)_ | 0.112 |  |  |  |  |  |  |

**Table S18.** Dependent variable: relative resilience (RRs), tree species: *Picea abies* from Poland

| Random effects | Name | Variance | *SD* |  |  |  |  |
| --- | --- | --- | --- | --- | --- | --- | --- |
| Plot | Intercept | 0.020 | 0.141 |  |  |  |  |
| Residual |  | 0.219 | 0.468 |  |  |  |  |
| Fixed effects |  |  |  |  |  |  |  |
| Variable | **β** | ***SE*** | ***t-value*** | ***df*** | ***P-value*** | **95% CI** | |
| Species richness | 0.037 | 0.049 | 0.752 | 17.607 | 0.462 | -0.066 | 0.139 |
| Live crown ratio | **0.146** | **0.066** | **2.202** | **98.244** | **0.030** | **0.014** | **0.278** |
| Age | -0.082 | 0.094 | -0.873 | 90.927 | 0.385 | -0.269 | 0.105 |
| Height | **-0.163** | **0.071** | **-2.300** | **103.082** | **0.023** | **-0.304** | **-0.022** |
| PreDr_ growth | -0.001 | 0.061 | -0.015 | 108.164 | 0.988 | -0.123 | 0.121 |
| Intercept | **-0.495** | **0.094** | **-5.287** | **53.172** | **0.000** | **-0.683** | **-0.307** |
| R^2^ _LMM (C)_ | 0.184 |  |  |  |  |  |  |
| R^2^ _LMM (M)_ | 0.109 |  |  |  |  |  |  |

**Table S19.** Dependent variable: recovery (Rc), tree species: *Picea abies* from Poland

| Random effects | Name | Variance | *SD* |  |  |  |  |
| --- | --- | --- | --- | --- | --- | --- | --- |
| Plot | Intercept | 0.007 | 0.086 |  |  |  |  |
| Residual |  | 0.164 | 0.404 |  |  |  |  |
| Fixed effects |  |  |  |  |  |  |  |
| Variable | **β** | ***SE*** | ***t-value*** | ***df*** | ***P-value*** | **95% CI** | |
| Species richness | 0.020 | 0.039 | 0.511 | 18.014 | 0.615 | -0.061 | 0.101 |
| Live crown ratio | 0.064 | 0.056 | 1.148 | 93.423 | 0.254 | -0.047 | 0.175 |
| Age | -0.030 | 0.079 | -0.381 | 83.259 | 0.705 | -0.186 | 0.126 |
| Height | **-0.176** | **0.060** | **-2.945** | **97.988** | **0.004** | **-0.294** | **-0.057** |
| PreDr_growth | -0.016 | 0.052 | -0.308 | 109.908 | 0.758 | -0.120 | 0.088 |
| Intercept | **-0.437** | **0.077** | **-5.658** | **56.224** | **0.000** | **-0.591** | **-0.282** |
| R^2^ _LMM (C)_ | 0.140 |  |  |  |  |  |  |
| R^2^ _LMM (M)_ | 0.101 |  |  |  |  |  |  |

**Table S20.** Dependent variable: resistance (Rt), tree species: *Picea abies* from Poland

| Random effects | Name | Variance | *SD* |  |  |  |  |
| --- | --- | --- | --- | --- | --- | --- | --- |
| Plot | Intercept | 0.032 | 0.181 |  |  |  |  |
| Residual |  | 0.190 | 0.436 |  |  |  |  |
| Fixed effects |  |  |  |  |  |  |  |
| Variable | **β** | ***SE*** | ***t-value*** | ***df*** | ***P-value*** | **95% CI** | |
| Species richness | -0.083 | 0.051 | -1.609 | 16.850 | 0.126 | -0.191 | 0.026 |
| Live crown ratio | -0.109 | 0.064 | -1.707 | 104.277 | 0.091 | -0.237 | 0.018 |
| Age | 0.006 | 0.091 | 0.061 | 100.222 | 0.952 | -0.175 | 0.186 |
| Height | **0.163** | **0.068** | **2.391** | **108.169** | **0.019** | **0.028** | **0.298** |
| PreDr_growth | -0.046 | 0.058 | -0.796 | 104.952 | 0.428 | -0.161 | 0.069 |
| Intercept | **-0.333** | **0.094** | **-3.565** | **47.700** | **0.001** | **-0.522** | **-0.145** |
| R^2^ _LMM (C)_ | 0.251 |  |  |  |  |  |  |
| R^2^ _LMM (M)_ | 0.122 |  |  |  |  |  |  |

**Table S21.** Dependent variable: resilience (Rs), tree species: *Picea abies* from Romania

| Random effects | Name | Variance | *SD* |  |  |  |  |
| --- | --- | --- | --- | --- | --- | --- | --- |
| Plot | Intercept | 0.000 | 0.000 |  |  |  |  |
| Residual |  | 0.790 | 0.889 |  |  |  |  |
| Fixed effects |  |  |  |  |  |  |  |
| Variable | **β** | ***SE*** | ***t-value*** | ***df*** | ***P-value*** | **95% CI** | |
| Species richness | -0.137 | 0.108 | -1.262 | 91.000 | 0.210 | -0.352 | 0.078 |
| Live crown ratio | 0.142 | 0.090 | 1.576 | 91.000 | 0.118 | -0.037 | 0.322 |
| Age | 0.147 | 0.181 | 0.816 | 91.000 | 0.417 | -0.211 | 0.506 |
| Height | **-0.270** | **0.098** | **-2.757** | **91.000** | **0.007** | **-0.464** | **-0.075** |
| PreDr_ growth | 0.061 | 0.205 | 0.299 | 91.000 | 0.765 | -0.346 | 0.468 |
| Intercept | **0.623** | **0.133** | **4.670** | **91.000** | **0.000** | **0.358** | **0.888** |
| R^2^ _LMM (C)_ | NA |  |  |  |  |  |  |
| R^2^ _LMM (M)_ | 0.098 |  |  |  |  |  |  |

**Table S22.** Dependent variable: resistance (Rt), tree species: *Picea abies* from Romania

| Random effects | Name | Variance | *SD* |  |  |  |  |
| --- | --- | --- | --- | --- | --- | --- | --- |
| Plot | Intercept | 0.068 | 0.260 |  |  |  |  |
| Residual |  | 0.465 | 0.681 |  |  |  |  |
| Fixed effects |  |  |  |  |  |  |  |
| Variable | **β** | ***SE*** | ***t-value*** | ***df*** | ***P-value*** | **95% CI** |  |
| Species richness | -0.087 | 0.122 | -0.718 | 9.244 | 0.491 | -0.361 | 0.187 |
| Live crown ratio | 0.000 | 0.072 | -0.002 | 90.168 | 0.999 | -0.143 | 0.143 |
| Age | 0.000 | 0.149 | -0.002 | 88.823 | 0.999 | -0.297 | 0.296 |
| Height | **-0.170** | **0.081** | **-2.096** | **88.296** | **0.039** | **-0.331** | **-0.009** |
| PreDr_growth | -0.035 | 0.164 | -0.211 | 90.113 | 0.834 | -0.361 | 0.292 |
| Intercept | -0.020 | 0.130 | -0.158 | 20.517 | 0.876 | -0.290 | 0.249 |
| R^2^ _LMM (C)_ | 0.184 |  |  |  |  |  |  |
| R^2^ _LMM (M)_ | 0.065 |  |  |  |  |  |  |

**Table S23.** Dependent variable: relative resilience (RRs), tree species: *Pinus sylvestris* from Spain

| Random effects | Name | Variance | *SD* |  |  |  |  |
| --- | --- | --- | --- | --- | --- | --- | --- |
| Plot | Intercept | 0.003 | 0.058 |  |  |  |  |
| Residual |  | 0.135 | 0.367 |  |  |  |  |
| Fixed effects |  |  |  |  |  |  |  |
| Variable | **β** | ***SE*** | ***t-value*** | ***df*** | ***P-value*** | **95% CI** | |
| Slope | 0.080 | 0.057 | 1.390 | 15.868 | 0.184 | -0.042 | 0.202 |
| Species richness | **0.203** | **0.050** | **4.068** | **15.045** | **0.001** | **0.096** | **0.309** |
| Crown live ratio | -0.020 | 0.044 | -0.455 | 83.337 | 0.650 | -0.109 | 0.068 |
| BAT subplot | 0.044 | 0.045 | 0.970 | 91.929 | 0.335 | -0.046 | 0.133 |
| Age | -0.033 | 0.052 | -0.642 | 86.229 | 0.523 | -0.137 | 0.070 |
| Height | 0.085 | 0.127 | 0.673 | 41.494 | 0.505 | -0.171 | 0.342 |
| PreDr_ growth | 0.096 | 0.232 | 0.414 | 91.689 | 0.680 | -0.365 | 0.557 |
| Intercept | **0.401** | **0.139** | **2.877** | **40.463** | **0.006** | **0.119** | **0.683** |
| R^2^ _LMM (C)_ | 0.310 |  |  |  |  |  |  |
| R^2^ _LMM (M)_ | 0.293 |  |  |  |  |  |  |

**Table S24.** Dependent variable: recovery (Rc), tree species: *Pinus sylvestris* from Spain

| Random effects | Name | Variance | *SD* |  |  |  |  |
| --- | --- | --- | --- | --- | --- | --- | --- |
| Plot | Intercept | 1.588 | 1.260 |  |  |  |  |
| Residual |  | 4.427 | 2.104 |  |  |  |  |
| Fixed effects |  |  |  |  |  |  |  |
| Variable | **β** | ***SE*** | ***t-value*** | ***df*** | ***P-value*** | **95% CI** | |
| Slope | 0.556 | 0.523 | 1.062 | 11.450 | 0.310 | -0.591 | 1.702 |
| Species richness | **1.140** | **0.455** | **2.505** | **11.627** | **0.028** | **0.145** | **2.135** |
| Crown live ratio | -0.336 | 0.280 | -1.201 | 91.316 | 0.233 | -0.893 | 0.220 |
| BAT subplot | 0.209 | 0.273 | 0.765 | 91.023 | 0.447 | -0.334 | 0.752 |
| Age | 0.086 | 0.321 | 0.267 | 91.291 | 0.790 | -0.551 | 0.723 |
| Height | 0.884 | 0.857 | 1.031 | 85.447 | 0.306 | -0.821 | 2.588 |
| PreDr_growth | 0.517 | 1.399 | 0.370 | 88.923 | 0.712 | -2.263 | 3.298 |
| Intercept | **1.984** | **0.976** | **2.034** | **61.059** | **0.046** | **0.033** | **3.935** |
| R^2^ _LMM (C)_ | 0.409 |  |  |  |  |  |  |
| R^2^ _LMM (M)_ | 0.197 |  |  |  |  |  |  |

**Table S25.** Dependent variable: resistance (Rt), tree species: *Pinus sylvestris* from Spain

| Random effects | Name | Variance | *SD* |  |  |  |  |
| --- | --- | --- | --- | --- | --- | --- | --- |
| Plot | Intercept | 0.031 | 0.170 |  |  |  |  |
| Residual |  | 0.108 | 0.329 |  |  |  |  |
| Fixed effects |  |  |  |  |  |  |  |
| Variable | **β** | ***SE*** | ***t-value*** | ***df*** | ***P-value*** | **95% CI** | |
| Slope | -0.036 | 0.076 | -0.472 | 12.549 | 0.645 | -0.201 | 0.129 |
| Species richness | **-0.203** | **0.066** | **-3.065** | **12.668** | **0.009** | **-0.347** | **-0.06** |
| Crown live ratio | 0.077 | 0.043 | 1.788 | 90.707 | 0.077 | -0.009 | 0.164 |
| BAT subplot | -0.04 | 0.043 | -0.931 | 91.41 | 0.354 | -0.124 | 0.045 |
| Age | 0.001 | 0.05 | 0.022 | 91.799 | 0.982 | -0.098 | 0.1 |
| Height | -0.08 | 0.132 | -0.604 | 82.242 | 0.547 | -0.342 | 0.183 |
| PreDr_growth | -0.333 | 0.218 | -1.531 | 89.995 | 0.129 | -0.766 | 0.099 |
| Intercept | **-0.802** | **0.149** | **-5.396** | **61.856** | **0** | **-1.1** | **-0.505** |
| R^2^ _LMM (C)_ | 0.410 |  |  |  |  |  |  |
| R^2^ _LMM (M)_ | 0.242 |  |  |  |  |  |  |

**Table S26.** Dependent variable: resilience (Rs), tree species: *Quercus robur* from Poland.

| Random effects | Name | Variance | *SD* |  |  |  |  |
| --- | --- | --- | --- | --- | --- | --- | --- |
| Plot | Intercept | 0.000 | 0.000 |  |  |  |  |
| Residual |  | 0.164 | 0.405 |  |  |  |  |
| Fixed effects |  |  |  |  |  |  |  |
| Variable | **β** | ***SE*** | ***t-value*** | ***df*** | ***P-value*** | **95% CI** | |
| Species richness | **0.100** | **0.050** | **1.995** | **87** | **0.049** | **0.000** | **0.200** |
| Crown live ratio | -0.115 | 0.084 | -1.373 | 87 | 0.173 | -0.282 | 0.052 |
| BAT subplot | -0.078 | 0.054 | -1.443 | 87 | 0.153 | -0.186 | 0.030 |
| Age | **0.090** | **0.038** | **2.371** | **87** | **0.020** | **0.015** | **0.166** |
| Height | -0.170 | 0.110 | -1.546 | 87 | 0.126 | -0.388 | 0.048 |
| PreDr_growth | **-0.849** | **0.319** | **-2.658** | **87** | **0.009** | **-1.483** | **-0.214** |
| Intercept | **-0.558** | **0.137** | **-4.060** | **87** | **0.000** | **-0.831** | **-0.285** |
| R^2^ _LMM (C)_ | NA |  |  |  |  |  |  |
| R^2^ _LMM (M)_ | 0.214 |  |  |  |  |  |  |
